# Mechanisms of translation repression by the EIF4E1-4EIP cap-binding complex of *Trypanosoma brucei*: potential roles of the NOT complex and a terminal uridylyl transferase

**DOI:** 10.1101/2021.05.19.444837

**Authors:** Franziska Falk, Kevin Kamanyi Marucha, Christine Clayton

## Abstract

Most transcription in *Trypanosoma brucei* is constitutive and polycistronic. Consequently, the parasite relies on post-transcriptional mechanisms, especially affecting translation initiation and mRNA decay, to control gene expression both at steady-state and for adaptation to different environments. The parasite has six isoforms of the cap-binding protein EIF4E as well as five EIF4Gs. EIF4E1 does not bind to any EIF4G, instead being associated with a 4E-binding protein, 4EIP. 4EIP represses translation and reduces the stability of a reporter mRNA when artificially tethered to the 3’-UTR, whether or not EIF4E1 is present. 4EIP is essential during the transition from the mammalian bloodstream form to the procyclic form that lives in the Tsetse vector. In contrast, EIF4E1 is dispensable during differentiation, but is required for establishment of growing procyclic forms. There are two competing models for EIF4E1 function: either EIF4E1 has translation initiation activity that is inhibited by 4EIP, or EIF4E1 acts only as an inhibitor. We here provide evidence for the second hypothesis. Even in the complete absence of 4EIP, EIF4E1 showed no detectable association with other translation initiation factors, and 4EIP loss caused no detectable change in 4E1-associated mRNAs. We found that 4EIP stabilises EIF4E1, probably through co-translational complex assembly, and that 4EIP directly recruits the cytosolic terminal uridylyl transferase TUT3 to EIF4E1/4EIP complexes. There was, however, no evidence that TUT3 is essential for 4EIP function; instead, some evidence implicated the NOT deadenylase complex.

## INTRODUCTION

The rates at which translation initiation and elongation occur determine the amount of protein that is synthesized from a particular mRNA (Firczuk et al. 2013). Cap-binding proteins exert essential roles during the initiation process. Classically, the cap is bound by an eIF4E protein, which in turn recruits an eIF4G protein. eIF4G serves as a scaffold for further association with the helicase eIF4A and the 43S complex, which consists of the small ribosomal subunit and several initiation factors. Once the latter has identified the first start codon through scanning, the large ribosomal subunit is recruited, allowing translation to proceed (Pestova et al. 2001). Most eukaryotic species studied possess more than one eIF4E homolog, the activities of which can either be restricted to a certain subset of mRNAs, or affect bulk translation (Hernandez et al. 2012).

Translation can be regulated through several mechanisms, many of which act upon the initiation process. In animal cells, binding of 4E-binding proteins (4E-BPs) and eIF4G proteins to eIF4Es is mutually exclusive, with the 4E-BP/eIF4E interaction leading to a blockade of productive initiation complex assembly (Kamenska et al. 2014b). Binding of 4E-BPs to eIF4Es is mediated through a YXXXXLØ motif near the 4E-BP N-terminus, which is shared with eIF4Gs (Mader et al. 1995). In contrast, eIF4E-like cap-binding proteins of the 4EHP-type do not bind to eIF4G proteins and have suppressive functions. They are exemplified by 4EHP/eIF4E2, d4EHP/EIF4E-8, and nCBP in mammals, *Drosophila*, and *Arabidopsis*, respectively (Rom et al. 1998; Hernandez et al. 2005; Kropiwnicka et al. 2015). Their suppressive functions appear to be mediated through their association with partner proteins, such as Bicoid or GIGYF2, which independently display RNA-binding activities (Cho et al. 2005; Morita et al. 2012; Fu et al. 2016; Peter et al. 2017; Amaya Ramirez et al. 2018). Association of mRNAs with either 4E-BP or 4EHP and their binding partners results in the recruitment of degradation machineries, such as deadenylation complexes, leading to reduced mRNA stability as well as reduced translation (Igreja and Izaurralde 2011; Amaya Ramirez et al. 2018). The 4EHP/GIGYF2 complex was recently implicated in co-translational destruction of mRNAs after ribosome pausing (Hickey et al. 2020; Juszkiewicz et al. 2020; Weber et al. 2020).

*Trypanosoma brucei* are unicellular eukaryotic parasites, members of the order Kinetoplastida which also includes *Leishmania* species. Kinetoplastids regulate gene expression primarily through post-transcriptional mechanisms, as transcription by RNA polymerase II is constitutive and polycistronic (Clayton 2016). However, they are faced with a high demand for plasticity in terms of gene expression profiles, as nutrient sources, ambient temperature, host defense mechanisms, and other factors differ vastly between the hosts encountered throughout the life cycle. For *Trypanosoma brucei* in the bloodstream of a mammalian host, glucose serves as the main source for ATP production, and evasion of the immune response by antigenic variation of surface proteins is required for survival (Vickerman 1978; Barbour and Restrepo 2000; Horn 2014). As trypanosome cell densities reach a critical level, development of long slender bloodstream forms into stumpy forms is induced. Growth arrest and low overall mRNA levels and translation rates are hallmark features of the stumpy stage, but expression of a selected set of proteins is strongly activated, including the surface protein PAD1 and some proteins of mitochondrial energy metabolism (Pays et al. 1993; Brecht and Parsons 1998; Dean et al. 2009). The stumpy form is primed for differentiation into procyclic forms upon uptake by a tsetse fly through a blood meal. Procyclic forms express EP and GPEET procyclins on their surface, and amino acids serve as main sources for energy production (Bringaud et al. 2006; Mantilla et al. 2017).

In accordance with their need for adaptive translation regulation, kinetoplastid parasites are equipped with six eIF4E isoforms (EIF4E1-6), as well as five eIF4Gs (EIF4G1-5), the individual roles of which have been the subject of numerous studies (Freire et al. 2017). EIF4Es 3-6 have EIF4G partners and are probably all active in translation initiation, while the role of EIF4E2 is unclear (Freire et al. 2018; Baron et al. 2021). Much of the evidence concerning EIF4E1 comes from *Leishmania*, which grow intracellularly as amastigotes within vertebrates, and as promastigotes within the sandfly vector. *Leishmania* EIF4E1 does not interact with any of the EIF4Gs, instead associating with a protein called 4E-interacting protein, 4EIP, which has the canonical EIF4E binding motif YXXXXLØ at its N-terminus (Zinoviev et al. 2011). A second EIF4E-binding protein, 4EBP2, was also recently found to associate with either EIF4E1 or EIF4E3 (Tupperwar et al. 2020). Leishmania promastigotes that completely lacked EIF4E1 showed altered morphology, with impaired growth and reduced overall translation and metabolism; the proteome changes in the mutant cells reflected their greatly reduced flagellar length. The ability of these cells to infect, and multiply within, macrophages was reduced, but not completely eliminated (Tupperwar et al. 2019).

After *Leishmania* EIF4E1 purification, some general translation initiation factors, including EIF3 and EIF2 subunits, were detected. In addition, tagged *Leishmania* EIF3A is able to pull down EIF4E1 *in vitro*, and a yeast 2-hybrid interaction between EIF3A and EIF4E1 was demonstrated (Meleppattu et al. 2015), suggesting that *Leishmania* EIF4E1 might act in translation initiation by direct EIF3 recruitment. Structural and *in vitro* studies characterised EIF4E1 binding to an N-terminal fragment of 4EIP, and showed that 4EIP inhibits EIF4E1 binding to the cap (Meleppattu et al. 2018). These results would all be consistent with a model in which *Leishmania* EIF4E1 is an EIF4G-independent translation initiation factor whose activity is inhibited by 4EIP and perhaps also by 4EIP2.

Results with *T. brucei* have confirmed the interaction between EIF4E1 and 4EIP, but have also suggested that EIF4E1 and 4EIP might have some independent activities. When tethered via a λN peptide to a boxB-bearing reporter mRNA, *T. brucei* 4EIP strongly represses reporter expression whether or not EIF4E1 is present (Erben et al. 2014; Terrao et al. 2018). This could mean that the tethering is merely replacing EIF4E1 in recruiting 4EIP to RNA, since similar activities have also been shown for metazoan 4E-BPs (Igreja and Izaurralde 2011; Kamenska et al. 2014a). However we showed that 4EIP, like GIGYF2, is directly cross-linkable to mRNAs (Lueong et al. 2016), which might allow it to function independently.

Although bloodstream forms lacking either EIF4E1 or 4EIP have only a mild growth defect, 4EIP is essential for full differentiation to the stumpy form; without 4EIP, the translational repression that is intrinsic to this process is delayed and the cells are no longer able to differentiate fully to procyclic forms (Terrao et al. 2018). Notably, a truncated form of 4EIP that cannot bind to EIF4E1 can restore stumpy formation in the 4EIP-deficient cells (Terrao et al. 2018). Meanwhile, initial differentiation of EIF4E1-deficient cells proceeds normally. These observations suggest a function of 4EIP that is independent of EIF4E1. Although tethering of EIF4E1 causes repression when 4EIP is present, in the absence of 4EIP tethering of EIF4E1 has no effect at all. One observation, however, suggests that EIF4E1 has an independent function; unlike 4EIP, we were unable to obtain procyclic trypanosomes lacking EIF4E1 (Terrao et al. 2018). Apart from this, Mabille et al. highlighted that 4EIP is essential for *in vivo* infectivity in the vertebrate host, as evidenced by a failure to detect 4EIP KO parasites after one week of infection (Mabille et al. 2021); immunosuppressive treatment could not rescue the defect.

This paper also concerns a *T. brucei* terminal uridylyl transferase. In mammalian cells and *Schizosaccharomyces pombe*, addition of uridine residues to the 3’-ends of RNAs makes them substrates of the 3’-5’ exoribonuclease Dis3L2 (Viegas et al. 2015), with subsequent degradation by the exosome. Substrates include U6 snRNA in the nucleus, and miRNAs and the non-polyadenylated histone mRNAs in the cytosol (Trippe et al. 2006; Lackey et al. 2016). *T. brucei* has five proteins with strong similarity to terminal uridylyl transferases (Aphasizhev et al. 2004). Three of them are implicated in mitochondrial RNA editing (Aphasizhev and Aphasizheva 2011). One of the remaining ones, TUT3 (Tb927.10.7310), is a highly processive terminal uridylyl transferase, lacks a mitochondrial targeting signal and, by cell fractionation, was not in the mitochondria of *Leishmania tarentolae* (Aphasizhev et al. 2004). Neither the level of the *TUT3* mRNA, nor its translation, are developmentally regulated in *T. brucei* (Jensen et al. 2014) and mass spectrometry results suggest that it has low abundance and is absent from mitochondria (Gunasekera et al. 2012; Dejung et al. 2016). A Dis3L2 homolog (Tb927.11.8290) is present, but the protein is concentrated in the nucleus (Fritz et al. 2015). In a high-throughput RNAi screen using the Lister 427 strain of *T. brucei*, which is unable to make stumpy forms or transform to growing procyclic forms, RNAi targeting either TUT3 or the Dis3L2 homologue had no effect on bloodstream form or procyclic form trypanosome survival (Alsford et al. 2011).

The main aim of the current study was to clarify the possible roles of EIF4E1 both with, and independent of 4EIP. We found no evidence that EIF4E1 can act as a translation initiation factor, but did discover an unexpected interaction between 4EIP and TUT3.

## RESULTS

### EIF4E1 is expressed in both bloodstream forms and procyclic forms, where it is stabilised by 4EIP

Analysis of EIF4E1 using specific antibody showed that it is expressed in both life cycle stages, with relative expression being slightly higher in the bloodstream form (**Fig. 1A**). All experiments were done in differentiation-competent EATRO1125-strain trypanosomes expressing the *Tn10* tet repressor. To follow EIF4E1 expression more easily in wild-type (WT) and 4EIP knockout (KO) backgrounds, we introduced a sequence encoding a PTP-tag into the genomic *EIF4E1* locus, joining the tag to the protein C-terminus (**Fig. 1B**). Concomitant KO of the second copy in procyclic forms gave no growth defect, confirming the function of the fusion protein. Both EIF4E1-PTP (**Fig. 1B**) and untagged EIF4E1 (**Fig. 1C, D**) showed reduced abundance in the absence of 4EIP, and expression could be partially and fully rescued in bloodstream forms and procyclic forms, respectively, upon ectopic expression of full-length 4EIP-myc (**Fig. 1C, D**). A deletion mutant of 4EIP lacking the N-terminal interaction motif for binding to EIF4E1 was unable to rescue EIF4E1 expression (**Fig. 1D**). These results suggest that interaction of 4EIP with EIF4E1 is required in order to maintain normal levels of EIF4E1.

**Fig. 1.**
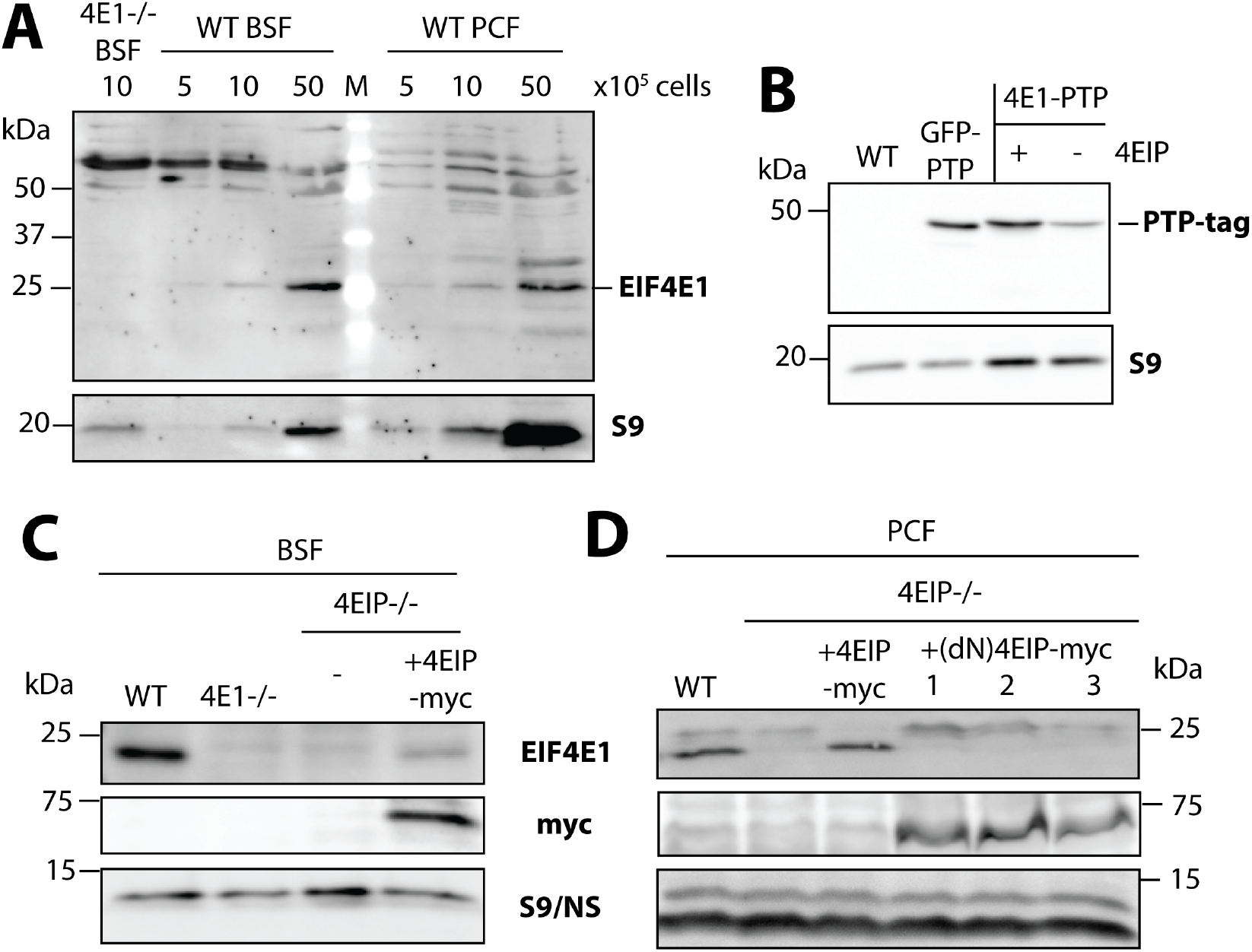
EIF4E1 expression and stability in bloodstream and procyclic *Trypanosoma brucei* parasites. **(A)** Bloodstream and procyclic forms (BSF and PCF, respectively) were collected at the numbers indicated and analysed for expression of EIF4E1 by western blotting with specific antibodies. **(B)** Western blot analysis of PTP-tagged versions of EIF4E1 (± 4EIP) and GFP, which were introduced into pleomorphic bloodstream forms of *T. brucei*. These could be switched to procyclic forms by *cis*-aconitate treatment (not shown). **(C)** Expression of EIF4E1 was analysed by western blotting in WT, EIF4E1- and 4EIP-deficient bloodstream forms, as well as in 4EIP-deficient cells reconstituted with full-length, myc-tagged 4EIP. **(D)** Expression of EIF4E1 was analysed in WT and 4EIP-deficient procyclic forms, as well as in 4EIP-deficient cells reconstituted with myc-tagged full-length or N-terminally deleted 4EIP. NS, non-specific control; S9, 40S ribosomal protein S9

*In situ* C-terminal tagging replaces the mRNA 3’-untranslated region, which is normally important for expression regulation; it is therefore possible that abnormal levels of total EIF4E1 are produced in the tagged cells. However, the requirement for 4EIP to maintain the protein will counteract such effects.

### EIF4E1 does not associate with general translation initiation factors in the absence of 4EIP

It was previously suggested that *Leishmania* EIF4E1 acts as a translation factor which is inhibited by 4EIP (Meleppattu et al. 2015). One piece of evidence was the *in vitro* and two-hybrid interaction between *Leishmania* EIF4E1 and EIF3A (Meleppattu et al. 2015). To find out whether this interaction occurs *in vivo* in *T. brucei*, we pulled down EIF4E1-PTP from cells expressing V5-tagged EIF3A. No association was detected, whether or not 4EIP was present (**Fig. S1A, B**). To get a broader understanding of EIF4E1 interactions, the protein binding partners of EIF4E1-PTP were analysed by quantitative mass spectrometry. This was done both in bloodstream forms, and in procyclic forms expressing only the tagged version. The pull-downs were done in the presence or absence of 4EIP. Notably, no general translation factors were enriched with EIF4E1-PTP, even when 4EIP was absent (**Fig. 2A-D**). *T. brucei* 4EIP2 (Tb927.10.11000) was enriched with EIF4E1 in both forms, whether 4EIP was present or not (**Fig. 2C, D**). In striking contrast, apart from 4EIP itself, the terminal uridylyl transferase TUT3 was present in EIF4E1 complexes only if 4EIP was present (**Fig 2A, B**). A possible association with components of the CAF1-NOT deadenylation complex was observed only for bloodstream forms (**Fig. 2B**).

**Fig. 2.**
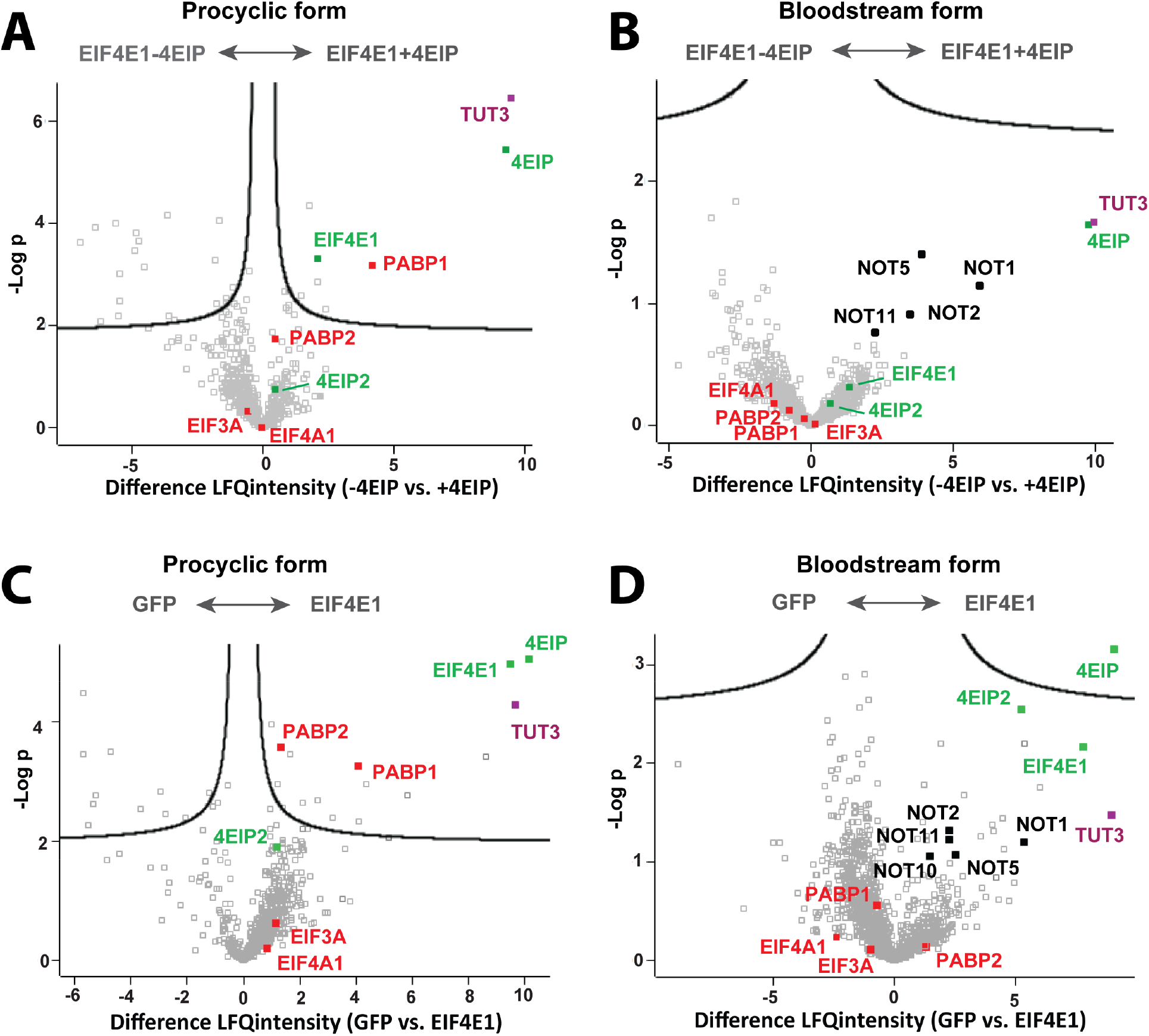
Protein binding partners of EIF4E1 in bloodstream and procyclic forms (BSFs and PCFs, respectively) of *Trypanosoma brucei*. **(A)** PTP-tagged EIF4E1 was pulled down from procyclic forms for comparison of bound proteins in WT and 4EIP knockout backgrounds using quantitative mass spectrometry. **(B)** PTP-tagged EIF4E1 was pulled down from bloodstream forms for comparison of bound proteins in WT and 4EIP knockout backgrounds using quantitative mass spectrometry. **(C)** PTP-tagged GFP and EIF4E1 were pulled down from procyclic forms for comparison of bound proteins using quantitative mass spectrometry. **(D)** PTP-tagged GFP and EIF4E1 were pulled down from bloodstream forms for comparison of bound proteins using quantitative mass spectrometry.

To confirm the interactions detected by mass spectrometry, co-immunoprecipitations were performed. TUT3 was reliably pulled down with 4EIP, and vice versa, whether RNase was added or not (**Fig. S2A, B**). On the other hand, the putative interaction with the NOT complex, judged by co-immunoprecipitation of the exonuclease CAF1, was weak (**Fig. S2C**) and not reproducible (**Fig. S2D**). Notably, we had previously obtained similar results concerning interactions of 4EIP in bloodstream forms: NOT1 was enriched with 4EIP by mass spectrometry, but no interaction was detected by co-immunoprecipitation (Terrao et al. 2018). These results suggest that 4EIP undergoes transient, low-stoichiometry interactions with the NOT complex.

### TUT3 is recruited by direct interaction with 4EIP

To evaluate whether the interaction between TUT3 and 4EIP was direct, yeast 2-hybrid assays were performed, further including the *T. brucei* homolog of Dis3L2. Yeast cells could grow on plates lacking adenine and histidine when both 4EIP and TUT3 were expressed; this was more pronounced with the combination of TUT3-AD (activation domain) and 4EIP-BD (DNA-binding domain) (**Fig. 3A, B**) than vice versa. The cells could also survive the selection process to some extent when 4EIP-AD and 4EIP-BD were expressed, suggesting self-interaction. The cells failed to grow with any of the other combinations tested, except for the positive control. The combined results therefore suggest that 4EIP directly recruits TUT3.

**Fig. 3.**
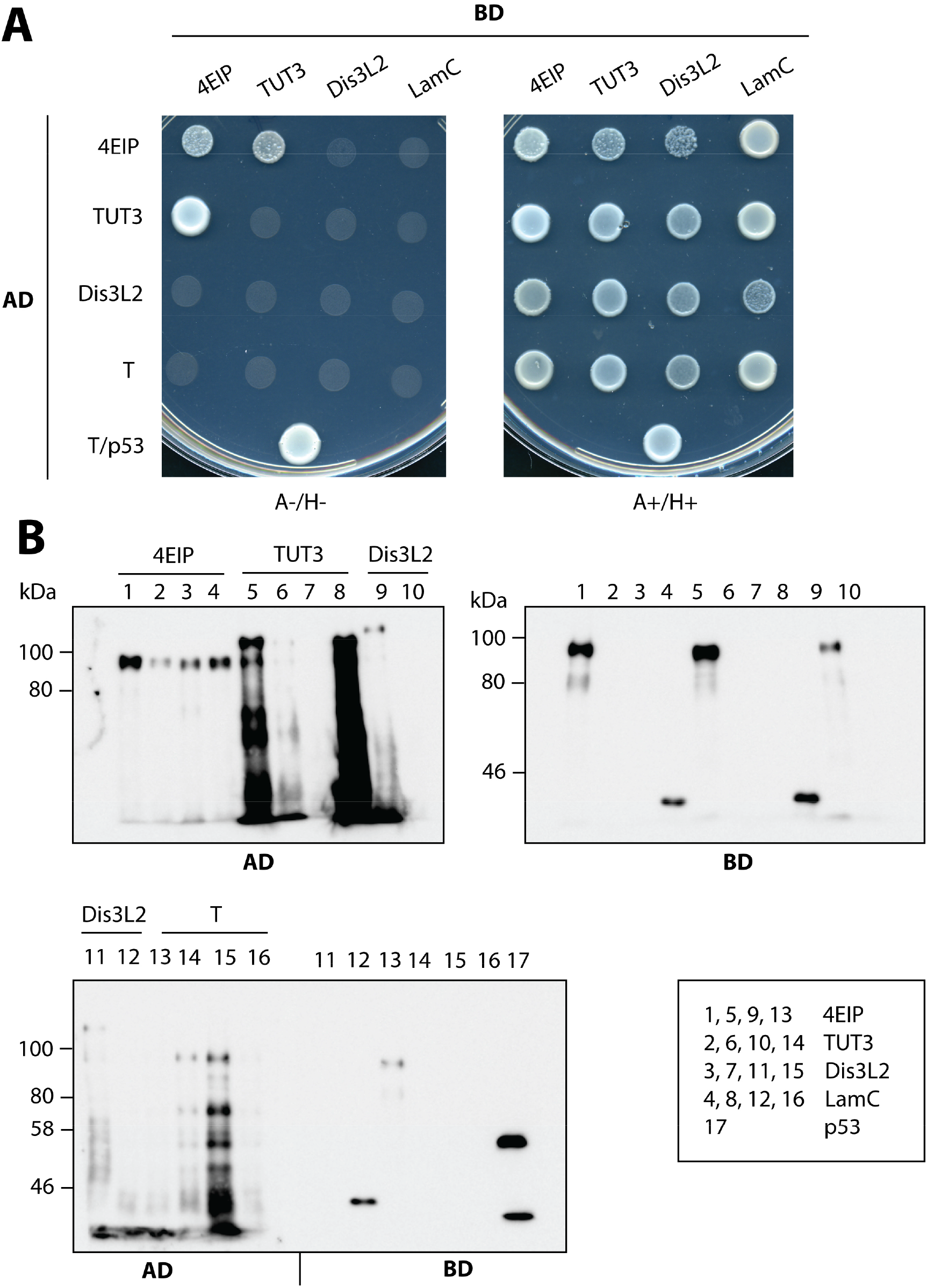
4EIP directly interacts with the terminal uridylyl transferase TUT3. **(A)** Yeast-2-hybrid assays for detection of direct interactions between 4EIP, TUT3, and the exonuclease Dis3L2, as indicated by growth on plates containing or lacking adenine and histidine (A+/H+ and A-/H-, respectively). The interaction between large T antigen (T) and p53 served as positive control. The combination of expression vectors encoding lamin C (LamC) and T served as negative control. **(B)** Western blotting for expression analysis of the proteins to be tested in the yeast-2-hybrid assay after transformation into yeast cells.

### TUT3 is a cytoplasmic enzyme that is essential in differentiation-competent bloodstream forms

TUT3 could cooperate with 4EIP only if it is in the cytosol. To confirm this, we expressed a C-terminally myc-tagged version. By immunofluorescence, TUT3-myc partially co-localised with both a cytosolic marker (**Fig. 4A**) and glycosomal aldolase (**Fig. 4B**), but upon digitonin fractionation, it was clearly in the cytosolic fraction (**Fig. 4C**).

**Fig. 4.**
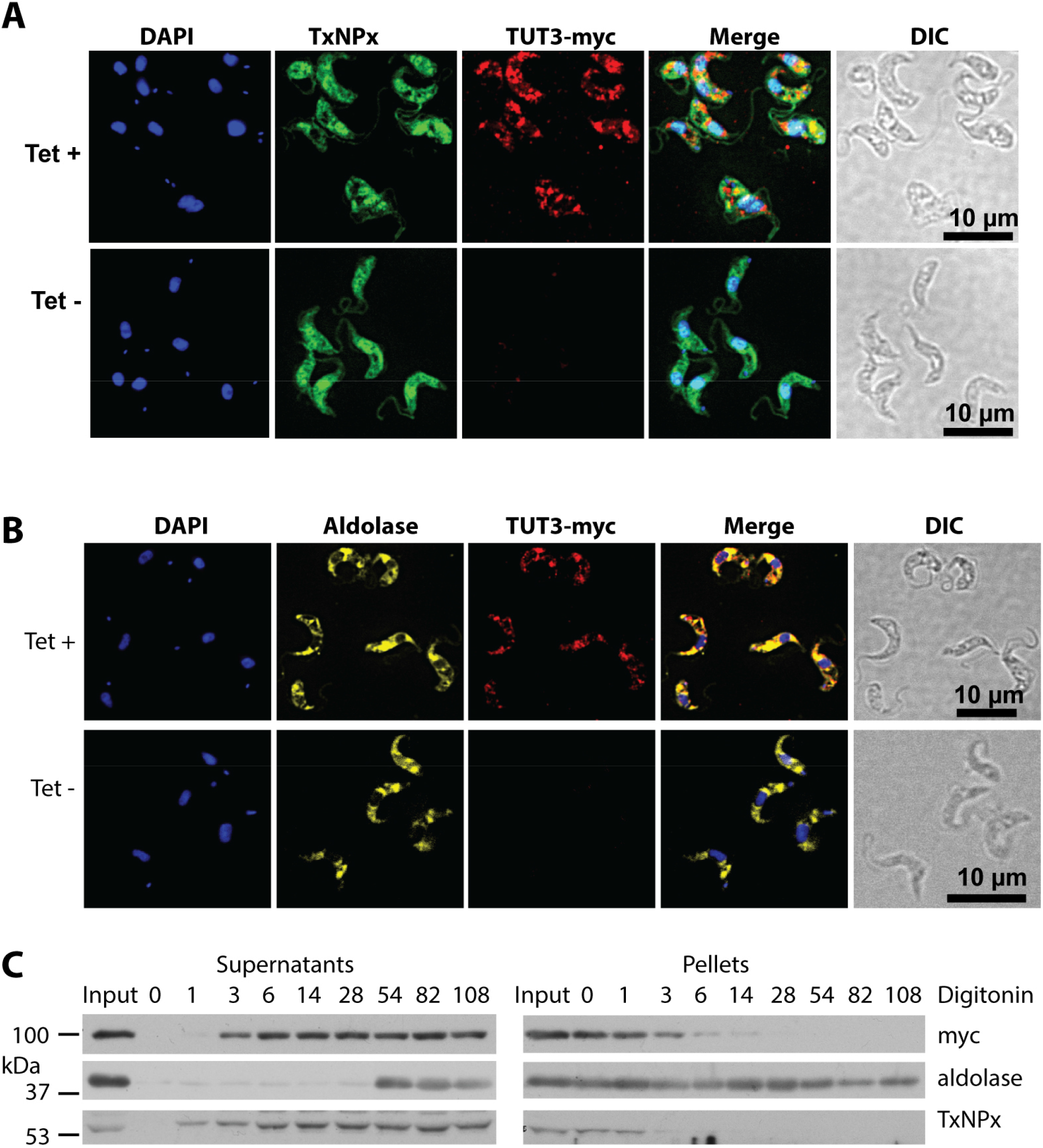
Location of C-terminally tagged TUT3. Bloodstream-form cells expressing tetracycline-inducible TUT3-myc were used. **(A, B)** Cells without tetracycline; for the upper panels, expression was induced for 24 h. In **(A)**, tryparedoxin peroxidase (TxNPx) was used as a cytosolic marker, and in **(B)**, aldolase as a glycosomal marker. DIC - differential interference contrast, DAPI - DNA stain. **(C)** Digitonin fractionation. The numbers on the lanes represent the mass of digitonin used per mass of protein (μg/mg) assuming that 5.4 × 10^6^ cells equal 100 mg of protein. Input is the total cell lysate.

To examine the function of TUT3 in differentiation-competent cells, we created bloodstream forms in which one copy of TUT3 bore a sequence encoding an N-terminal V5 tag. RNAi targeting TUT3 considerably reduced V5-TUT3 expression, but had no effect on growth of either bloodstream forms or procyclic forms (**Fig. S3A, B**). Since there is always residual expression after RNAi, we also attempted to knock out both *TUT3* genes. Multiple attempts to do this in the differentiation-competent cells failed. Interestingly, we did succeed in Lister 427 bloodstream forms, with no effect on growth (**Fig. S3C-E**). This result hints that TUT3 might be involved in aspects of gene expression control that are required only for the survival of fully differentiation-competent trypanosomes.

### Procyclic trypanosomes without 4EIP show reduced expression of GPEET mRNA

The results so far suggested that 4EIP might act by recruiting TUT3, resulting in 3’ uridylation and subsequent attack by DIS3L2 and then the exosome. The only uridylated mRNA described for *T. brucei* is that encoding the procyclin GPEET (Knusel and Roditi 2013). To find out whether TUT3 is involved in the regulation of GPEET expression, we induced differentiation of bloodstream forms with TUT3 RNAi by adding *cis*-aconitate and a shift in temperature from 37 °C to 27 °C. Expression of GPEET was induced in both WT and TUT3-V5 RNAi cells, with a certain delay in the case of the latter (**Fig. 5A, B**), while expression of EP procyclin was induced with normal kinetics (**Fig S4**). Since, however, RNAi has low efficacy during growth arrest, the results are difficult to interpret. We therefore took another approach. Adaptation of procyclic forms to an environment high in glucose was previously shown to lead to reduced GPEET expression (Vassella et al. 2004). To analyse whether disruption of EIF4E1/4EIP/TUT3 complexes could affect GPEET expression, we compared expression of GPEET on the surface of WT and 4EIP KO cell lines either in our standard low-glucose medium or after growth in high glucose for three weeks (**Fig. 5C**). Intriguingly, cells lacking 4EIP had strongly reduced GPEET levels compared to WT cells even without growth in high-glucose medium. Nonetheless, as for WT cells, the level of GPEET expression was further reduced in response to the glucose treatment (**Fig. 5C**).

**Fig. 5.**
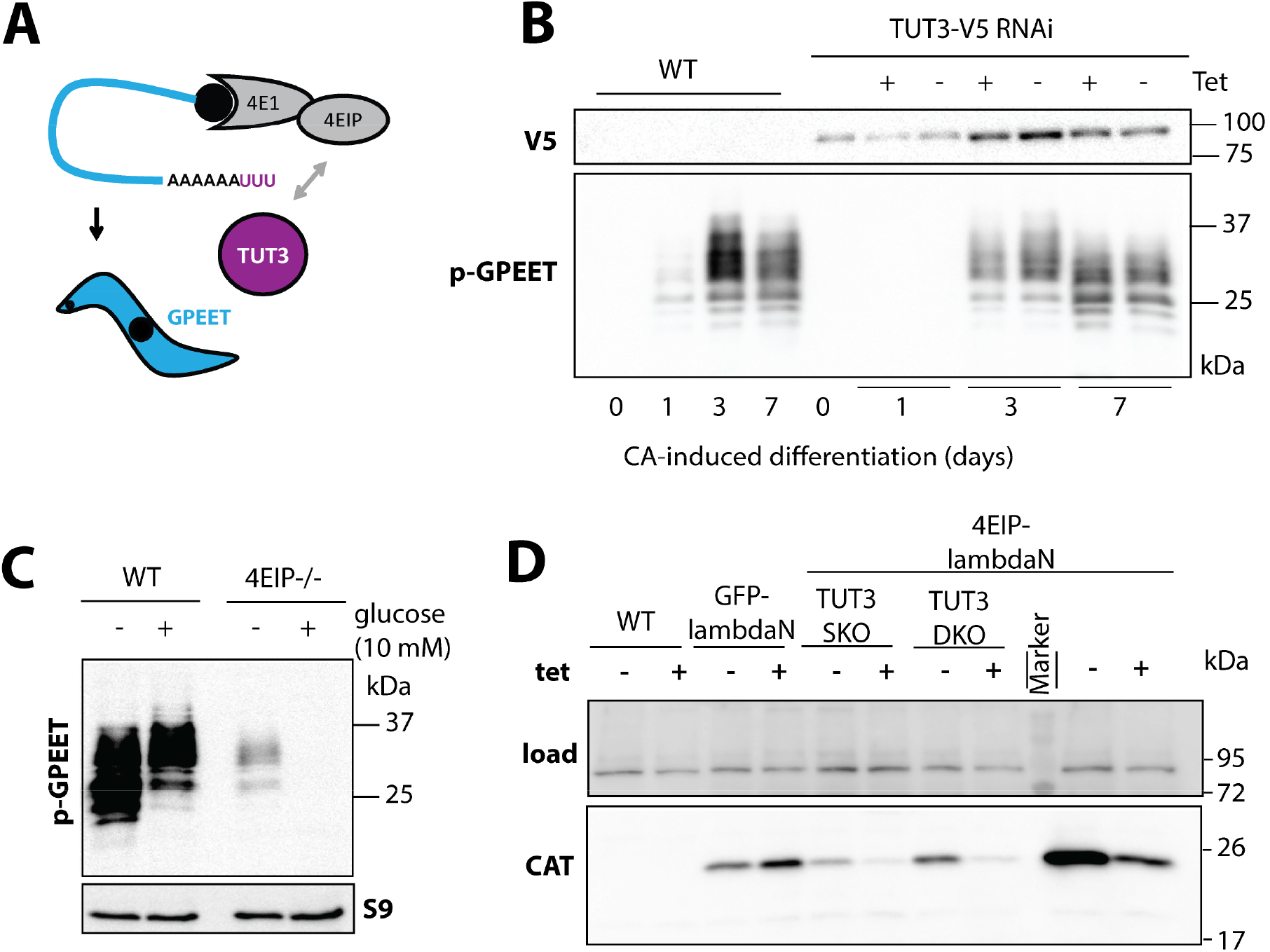
4E-interacting protein (4EIP) suppresses expression independently of TUT3. **(A)** Schematic representation of GPEET procyclin mRNA potentially serving as a substrate for TUT3-mediated uridylation upon recruitment by the EIF4E1/4EIP complex. **(B)** Analysis of phospho (p)-GPEET expression kinetics at the times indicated after *cis*-aconitate (CA)-induced differentiation in WT and TUT3-depleted cells. **(C)** Analysis of p-GPEET expression after adaptation of WT and 4EIP-deficient procyclic forms (PCFs) to 10 mM glucose. **(D)** Expression analysis of chloramphenicol acetyltransferase (CAT) by western blotting after tethering of 4EIP-lambdaN to the 3’UTR of the CAT mRNA through a box-B sequence in WT, TUT3 single-knockout, and TUT3 double-knockout backgrounds. As a control, GFP-lambdaN was tethered to the CAT mRNA in a WT background.

The results clearly showed that 4EIP is not required for glucose-induced loss of GPEET expression in procyclic forms. We therefore decided to check whether any uridylated *GPEET* mRNA was detectable in our procyclic cells. We examined GPEET mRNA 3’-ends in normal and glucose-adapted WT procyclic forms by cap removal, circularisation, selective amplification, and DNA sequencing. Poly(A)-tailed *GPEET* mRNAs were identified in 8 of 20 cloned products from low-glucose-grown cells, and only 1 of 20 from the high glucose cells; remaining clones were not *GPEET*. No uridylation was observed (Supplementary file **S6**). We also looked for potential targets of uridylation using four existing RNA-seq datasets from WT bloodstream forms and procyclic forms (**Fig. 6A**). We were able to identify several mRNAs containing at least 15 As followed by more than 5 Us at their 3’ ends. The levels of three mRNAs that were identified in at least three out of four data sets were compared in WT versus *TUT3* KO cells by RT-qPCR. Levels were similar between WT and *TUT3* KO cells for all three mRNAs analysed (**Fig. 6B**), ruling out a role for TUT3 in degradation of these mRNAs in Lister 427 bloodstream forms.

**Fig. 6.**
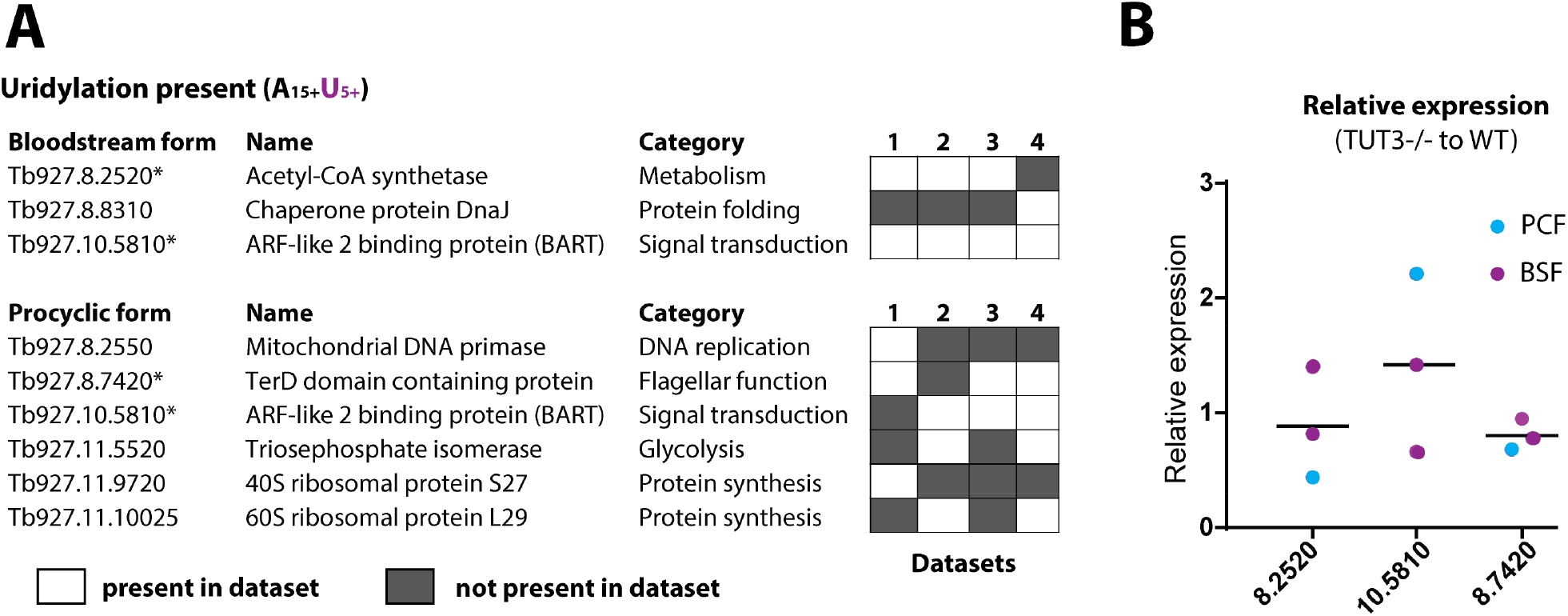
Identification of potential uridylation targets in *Trypanosoma brucei*. **(A)** *In silico* identification of potential 3’ uridylation targets by screening of RNA sequencing data from wild-type (WT) bloodstream forms (BSFs) and procyclic forms (PCFs) (four data sets each) for sequences containing at least 15 As followed by more than 5 Us. **(B)** Expression analysis of potential uridylation targets identified by the *in silico* analysis in **(A)** using quantitative reverse transcription PCR (qRT-PCR).

### TUT3 is not required for 4EIP-mediated repression when tethered to a reporter mRNA

We finally asked whether TUT3 was required in order for 4EIP to repress expression upon tethering to a reporter mRNA. For this, 4EIP fused to a λN peptide was artificially tethered to the 3’ UTR of the mRNA encoding chloramphenicol acetyltransferase (CAT) through a box-B sequence. This was done in WT, *TUT3* single-KO, and *TUT3* double-KO backgrounds. As shown in **Fig. 5D**, 4EIP could exert its repressive functions in absence of TUT3 in long slender bloodstream forms. Rather unexpectedly, tethering of TUT3 to the reporter had no effect on reporter expression (Supplementary **Fig. 4B**). This was surprising because we might at least have expected the tethered TUT3 to recruit 4EIP to mRNAs.

### The absence of 4EIP does not influence EIF4E1 mRNA association

There is *in vitro* evidence that 4EIP prevents EIF4E1 binding to the cap. If this is true *in vivo*, this should be evident from the EIF4E1-associated transcriptome. In the absence of 4EIP, binding of EIF4E1 to mRNAs should be increased. To investigate this, we performed pull-downs of EIF4E1-PTP in WT and *4EIP* KO backgrounds using procyclic forms, where EIF4E1 is essential. The bound protein was released with TEV protease (which cleaves within the tag), and RNAs in both the bound and unbound fractions were sequenced. Similar amounts of RNA were obtained in both purifications, but since only a single purification step was employed, with no replicates, no major conclusions can be drawn. We then compared the abundances of specific mRNAs in the bound and unbound fractions, in order to find out which mRNAs were specifically bound to EIF4E1. To our surprise, the results in the presence and absence of 4EIP were almost identical (**Fig. 7A**). There were two notable exceptions: the mRNAs encoding NOT1 and 4EIP, both of which were enriched about 60-fold in the presence of 4EIP. (Of course the 4EIP mRNA is not present in the absence of the gene; the few reads detected must be shared with other genes.) Since this was only a single experiment we confirmed the association in three additional independent pull-downs (**Fig. 7B**).

**Fig. 7.**
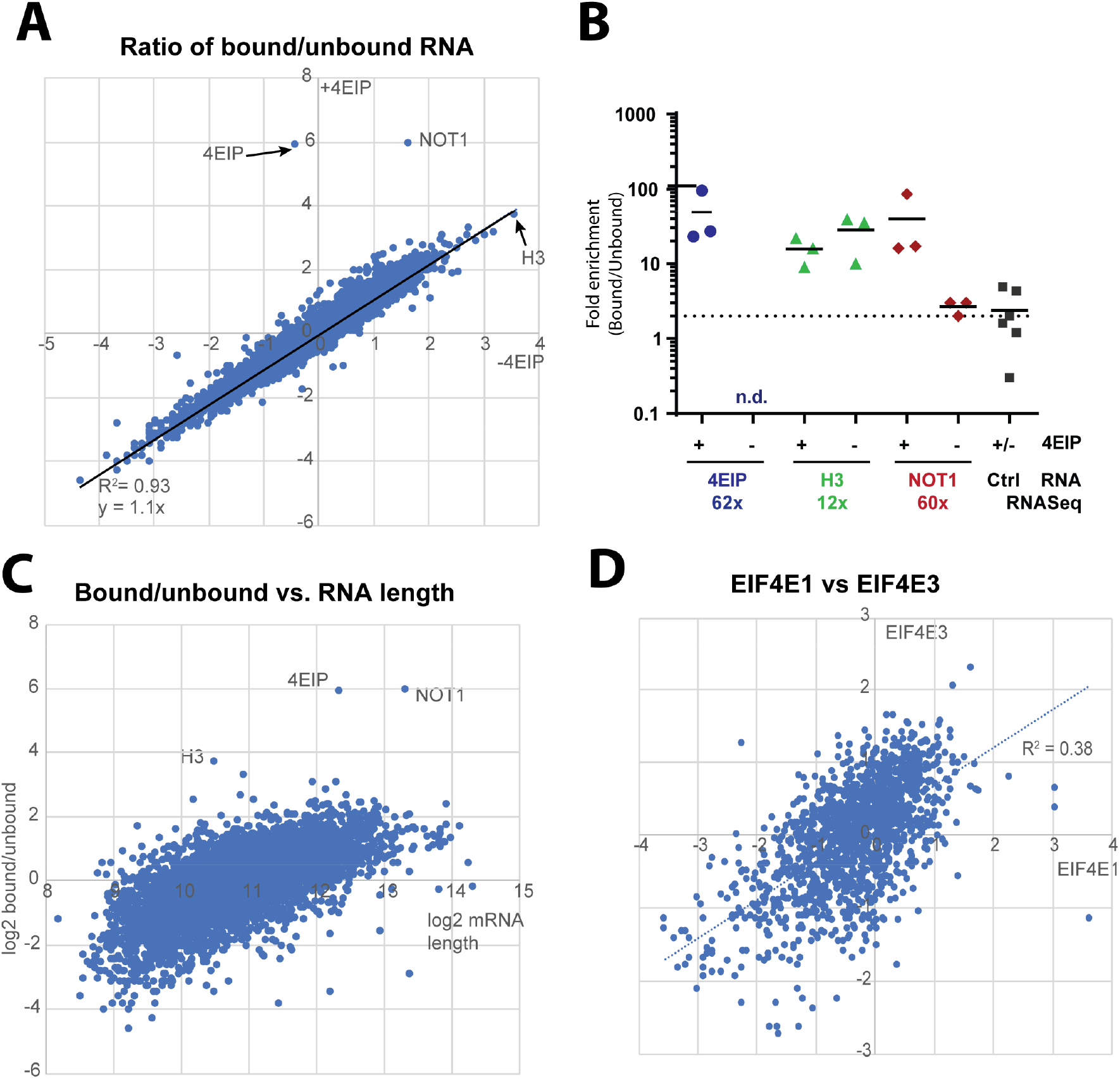
Identification of mRNAs bound to EIF4E1 ± 4E-interacting protein (4EIP) in *Trypanosoma brucei*. **(A)** The mRNAs pulled down with EIF4E1 (± 4EIP) from procyclic forms. The three most strongly enriched mRNAs are indicated. **(B)** Analysis of mRNAs found enriched with EIF4E1-PTP upon RNA sequencing using quantitative reverse transcription PCR (qRT-PCR) with specific primers. **(C)** Ratios of mRNA from EIF4E1-bound and -unbound fractions compared to mRNA length. **(D)** The mRNAs bound to EIF4E1 in procyclic forms were compared with those bound to EIF4E3 in bloodstream forms. The comparison was restricted to mRNAs that show no developmental regulation of abundance or ribosome occupancy

Apart from the two cases noted above, all mRNA interactions of EIF4E1 were similar whether 4EIP was present or not (**Fig. 7A**). We confirmed this for the histone H3 mRNA (**Fig. 7A, B**). The strong correlation seemed strange so we wondered about the specificity of the interactions. The specific association with 4EIP, NOT1 and histone H3 RNAs was superimposed on a background general preference for long mRNAs (**Fig. 7C**), which could result from relatively non-specific RNA-protein interactions (Erben et al. 2021). The preference for long RNAs also causes a bias towards mRNAs encoding RNA-binding proteins and protein kinases (Supplementary **Fig. S5A**).

To investigate this a bit more we compared the results with two other datasets: the mRNAs bound by 4EIP and by EIF4E3 in bloodstream forms. (The EIF4E3 data were generated for another project and will be presented in more detail elsewhere.) To avoid artefacts due to developmental regulation we restricted the analysis to mRNAs that show less than 1.5x regulation in both mRNA abundance (Fadda et al. 2014) and ribosome occupancy (Antwi et al. 2016). The results for EIF4E1 showed no correlation at all with those for 4EIP (Supplementary **Fig. S5B-D**), but did correlate with those for EIF4E3 (**Fig. 7D**, Supplementary **Fig. S5E, F**). Careful scrutiny suggested that less than ten mRNAs were specifically associated with EIF4E1; apart from those encoding 4EIP, Histone H3, and NOT1, they included Tb927.8.1670 (encoding the NEK14 protein kinase) and Tb927.6.1180 (a basal body protein). Overall the comparison confirmed our suspicion that background binding to (or trapping of) long mRNAs was obscuring all but the strongest specific EIF4E1-mRNA associations. To investigate this further it would be important to analyse translation in procyclic forms that have reduced levels of EIF4E1. We made numerous attempts to generate such cells using RNAi, but substantial amounts of EIF4E1 protein were always present after RNAi induction. Nevertheless, out results clearly provided no support for the idea that the *in vivo* function of 4EIP is to prevent binding of EIF4E1 to mRNA.

## DISCUSSION

The aim of this work was to clarify the functions of EIF4E1 both within and outside the complex with 4EIP. As a first step, tagged EIF4E1 was purified from differentiation-competent bloodstream form and procyclic forms and bound proteins were identified. Our results provided no evidence that 4EIP inhibits an *in vivo* translation initiation activity of EIF4E1; no specific associations of EIF4E1 with general translation factors were detected, even in the absence of 4EIP; and loss of 4EIP also did not result in enhanced association of EIF4E1 with mRNA. We did, however, have some hints regarding 4EIP function. We found clear evidence for association of 4EIP with the terminal uridylyl transferase TUT3, although its role is uncertain, and there were hints of weak association with the CAF1-NOT deadenylation complex. Cells lacking 4EIP also had decreased levels of EIF4E1 protein.

The protein 4EIP2 was associated with trypanosome EIF4E1 independently of 4EIP, suggesting binding to EIF4E1 as already indicated for *Leishmania* (Baron et al. 2021). Both N-and C-terminally tagged versions of *T. brucei* 4EIP2 are in the cytoplasm (Dean et al. 2017); high-throughput RNAi detected no defect for 4EIP2 depletion (Alsford et al. 2011), but the degree of depletion was of course not assessed. Pull-down results in *Leishmania* suggested that about a tenth of the EIF4E1 was 4EIP2-associated. Since 4EIP2 was not detected in our previous 4EIP pull-downs (Terrao et al. 2018), and the interaction most likely occurs via the same motif, 4EIP2 and 4EIP probably compete for EIF4E1 binding. It is possible that when 4EIP1 is absent, it is replaced by 4EIP2; this will depend on whether other EIF4Es are also competing for 4EIP2, and on the relative abundances of the different proteins.

Our mass spectrometry results yielded no evidence for association of EIF4E1 with general translation factors. The only representatives that were reproducibly found were EIF1 and EIF2, which were equally represented in the GFP control. Moreover, independent precipitation of EIF4E1-PTP, followed by western blotting, revealed no co-purification of EIF3A even if 4EIP was absent. Removal of 4EIP also (with two exceptions) had no effect on EIF4E1 mRNA association. In our previous tethering assays, EIF4E3, 4, 5 and 6 all were able to stimulate expression of the bound reporter RNA, but EIF4E1 had no effect (Erben et al. 2014). Although we cannot rule out a role for 4EIP2 in inhibiting EIF4E1 action, together these results provide no indication that EIF4E1 is able to drive translation initiation. Instead, we suggest that it supports repression by 4EIP.

Purification of EIF4E1-PTP resulted in extremely strong enrichment of *4EIP* mRNA. 4EIP interacts with EIF4E1 via the 4EIP N-terminus, so the *4EIP* mRNA might be connected to EIF4E1-PTP via the nascent 4EIP polypeptide, as is seen for some other protein complexes (Shiber et al. 2018). Since the EIF4E1-PTP was purified via a C-terminal tag, the result might imply that there is a pool of EIF4-PTP that is free for nascent 4EIP binding, and not bound by pre-existing 4EIP or 4EIP2. Conversely, levels of EIF4E1 were drastically reduced in cells lacking 4EIP, suggesting that the mutual interaction is required for EIF4E1 protein stability - a function that 4EIP2 cannot replace. The IBAQ values - although not reliable for absolute protein quantification - suggest roughly equimolar co-purification of 4EIP with EIF4E-PTP, and ten times less 4EIP2 (Supplementary table S1). Our results overall suggest that stabilisation of EIF4E1 by 4EIP may be initiated by co-translational folding of the two proteins.

Our results also yielded additional insights into the mechanism of action of 4EIP. Drosophila GIGYF2 - like 4EIP - is essential for repression by 4EHP, acting in part through recruitment of the NOT deadenylation complex (Ruscica et al. 2019). Purification of 4EIP or - in this paper, EIF4E1 with associated 4EIP - from bloodstream forms has reproducibly yielded results suggesting the association of small amounts of NOT complex. Although the association could not be confirmed by coimmunoprecipitation, this may be because the interaction is transient and involves only a small proportion of each complex. From the IBAQ values, there was at least ten times more 4EIP than NOT complex proteins in the pull-downs. In procyclic forms, the *NOT1* mRNA was also very strongly associated with EIF4E1, in a 4EIP-dependent fashion. We do not know what this result means: whether the mRNA is purified via the nascent polypeptide, or via direct RNA binding. In bloodstream forms, the *NOT1* mRNA was not enriched with 4EIP (Terrao et al. 2018); and in procyclic forms, there was no detectable co-purification of the NOT complex with EIF4E1 (**Fig. 2A, C**).

An alternative mechanism for 4EIP-mediated suppression would be through TUT3-mediated 3’-uridylation, followed by 3’ digestion by DIS3L2. Targets of uridylation are difficult to identify based on their (presumed) unstable nature. Our investigations concerning this were inconclusive. We were unable to confirm *GPEET* mRNA as a target of 3’-terminal uridylation, and the abundances of some other potential targets were unaffected in cells lacking TUT3. TUT3 also was not essential for the suppressive activities of 4EIP, at least when tethered to a reporter mRNA. The tethering experiments were, however, all done in differentiation-incompetent bloodstream forms, in which EIF4E1, 4EIP and TUT3 are not essential. We could not obtain differentiation-competent bloodstream forms, or procyclic forms, that completely lack TUT3, or lines with effective RNAi, so could not investigate this further. We hypothesise, however, that TUT3 might be needed to destabilise some mRNAs that promote differentiation in growing cells, and might also cooperate with the 4EIP-EIF4E1 complex.

## MATERIALS AND METHODS

### Trypanosome culture and differentiation

Monomorphic Lister 427 or pleomorphic AnTat 1.1 strains ectopically expressing the *Tet* repressor were used. bloodstream forms were cultured at 37 °C in HMI-9 medium (supplemented with 10 % (v/v) fetal calf serum (FCS), 1 % (v/v) penicillin/streptomycin solution (Labochem international, Germany), 15 μM L-cysteine, and 0.2 mM β-mercaptoethanol) in the presence of 5 % CO_2_ and 95 % humidity and their density was maintained below 1.0 × 10^6^ cells/mL unless stated otherwise. Procyclic forms were cultured at 27 °C in MEM-Pros media (supplemented with 10 % (v/v) FCS, 3.75 mg hemin, and 1 % (v/v) penicillin/streptomycin solution (Labochem international, Germany)), and their density was maintained between 5 × 10^5^ and 5 × 10^6^ cells/mL. Cell densities were determined using a Neubauer chamber.

To induce bloodstream form to procyclic form differentiation, cells grown at 1 × 10^6^ cells/mL in HMI-9 medium were incubated with 6 mM *cis*-aconiatete at 27 °C for 24 h. On the following day, the medium was exchanged for MEM and the cells were cultured at 27 °C afterwards.

### Immunofluorescence microscopy

Bloodstream forms at a density of 1.0 × 10^6^/mL were collected at 950 × *g* and washed with PBS, then fixed using 4 % para-formaldehyde for 20 min at RT. Fixed cells were left overnight at 4 °C to settle down on a chambered poly-L-lysine treated microscope slide. The cells were permeabilised by 0.2 % Triton X-100 for 20 min and subsequently washed three times using PBS in 5 min intervals. Blocking was done for 20 min using 0.5 % gelatin. The cells were incubated with primary antibodies (myc, 1:300; TxNPx, 1:1000; aldolase, 1:300) for 1 h and subjected to three more washing steps. The cells were incubated with secondary antibodies (1:500) coupled to fluorophores for 1 h, and washed twice before treatment with DAPI at 500 ng/mL for 15 min. The cells were washed twice and mounted using 90 % glycerol in PBS. Images were acquired using an Olympus IX81microscope and analysed with Olympus Xcellence or Image J software.

### Genetic manipulation of trypanosomes

Gene knockout was achieved by replacing the open reading frames (ORFs) of the target gene with ORFs encoding blasticidin and puromycin resistance genes, which integrate by homologous recombination. Double knock-out cells were selected by growing transfectants in both 5 μg/mL blasticidin and 0.2 μg/mL puromycin for at least two weeks before downstream analyses. Similarly, tagging with V5, myc or PTP tags was achieved by integration through homologous recombination in 5’- or 3’-UTRs for N- or C-terminal tagging of the protein of interest, followed by selection of transfectants with the respective drug. Of note, selection of pleomorphic clones was done using half the drug concentrations. The primers and plasmids used in this study can be found in Supplementary Table S3.

### DNA extraction

DNA was extracted from approximately 3 × 10^8^ cells as follows. The cell pellet was resuspended and the cells were lysed using 0.5 mL EB buffer (10 mM Tris-HCl pH 8.0, 10 mM NaCl, 10 mM EDTA). RNA was digested using 25 μg/mL of RNaseA (Sigma-Aldrich) at 37 °C for 30 min. Proteins were precipitated using 1.5 M ammonium acetate and centrifuging at 9,391 × *g* for 5 min. Isopropanol was added to the supernatant in a 1:1 ratio and centrifuged at 15,871 × *g* for 15 min at 4 °C. The pellet was washed with 75 % ethanol followed by centrifugation at 15,871 × *g* for 5 min and dried before resuspending in 50 μL TE buffer (10 mM Tris pH 7.5, 1 mM EDTA pH 8.0). PCR was done using Taq Polymerase according to the manufacturer’s instructions (New England Biolabs).

### Immunoprecipitations

For cell harvest, 2 × 10^9^ (for MS/RNA-IP) or 1 × 10^8^ (for WB) trypanosomes at a concentration of 5 × 10^5^ cells/mL (BSF) or 5 × 10^6^ cells/mL (PCF) were centrifuged at 3,000 rpm for 13 min at 4 °C. The pellet was resuspended in 50 mL of ice-cold PBS and centrifuged at 2,300 rpm for 8 min. After removal of the supernatant, the pellet was snap-frozen in liquid nitrogen and stored at −80 °C until further processing. All of the following steps were at 4 °C, unless stated otherwise. Cell pellets were thawed on ice and resuspended in 0.5 mL of lysis buffer (20 mM Tris [pH 7.5], 5 mM MgCl_2_, 1 mM DTT, 0.05 % IGEPAL, 100 U/mL RNasin, 10 μg/mL aprotinin, and 10 μg/mL leupeptin). For releasing protein contents, the cells were passaged 20× through a 21G×1½’’ needle and 20× through a 27G×¾ needle using a 1 mL syringe. In order to pellet the cell debris, samples were centrifuged at 10,000 × *g* for 15 min, and the supernatant was transferred to a fresh tube. The salt concentration was then adjusted to 150 mM KCl. Magnetic beads (Dynabeads™ M-280 Tosylactivated, Thermo Fisher Scientific) coupled to rabbit IgG were adjusted by three sequential washes with wash buffer (20 mM Tris [pH 7.5], 5 mM MgCl_2_, 1 mM DTT, 0.05% IGEPAL, 100 U/mL RNasin, 150 mM KCl). Depending on the cell number, 10-100 μL of the beads were then added to each sample. To allow binding, cell lysate and beads were incubated for 1-2 h at 4 °C while rotating (20 rpm).

The beads were boiled directly in Laemmli buffer for 10 min at 95 °C and analysed by western blotting (see below).

### Harvest for mass spectrometry of EIF4E1-associated proteins

Pull-downs were performed with 10^9^ bloodstream forms or procyclic forms according to the procedure described above. The beads were washed four times with wash buffer, after which bound proteins were released by TEV cleavage. For this, 20 μL of wash buffer and 1 μL of recombinant TEV protease (1 mg/mL) were incubated with the beads for 90 min at 20 °C. For removal of His-tagged TEV, IgG magnetic beads were concentrated on one side, the supernatant was transferred to a fresh tube, and 10 μL of equalization buffer (200 mM sodium phosphate, 600 mM sodium chloride, 0.1 % Tween-20, 60 mM imidazole, pH 8.5), as well as 30 μL of Ni-NTA-magnetic beads were added and incubated with the samples for 30 min at 20 °C while rotating. Ni-NTA magnetic beads were retained by a magnetic stand and the supernatant was collected and stored in Laemmli buffer at −80 °C.

Eluted proteins were separated on a 1.5 mm NuPAGE™ Novex™ 4-12 % Bis-Tris protein gel (Thermo Fisher Scientific) until the running front had migrated roughly 2 cm, after which the gel was stained with Coomassie blue and destained with destaining solution (10 % acetic acid, 50 % methanol in H2O). Three areas per lane were cut and analysed by Nanoflow LC-MS2 analysis with an Ultimate 3000 liquid chromatography system directly coupled to an Orbitrap Elite mass spectrometer (both Thermo-Fischer, Bremen, Germany). MS spectra (m/z 400–1600) were acquired in the Orbitrap at 60,000 (m/z 400) resolution. Fragmentation in CID cell was performed for up to 10 precursors. MS2 spectra were acquired at rapid scan rate. Raw files were processed using MaxQuant (version 1.5.3.30; J. Cox, M. Mann, Nat Biotechnol 2008, 26, 1367) for peptide identification and quantification. MS2 spectra were searched against the TriTrypDB-8.1TREU927-AnnotatedProteins-1 database (containing 11567 sequences). Data were analysed quantitatively and plotted using Perseus software (Version 1.6.6.0). Raw data are available in the PRIDE database with the accession number PXD025913.

### Harvest for isolation or EIF4E1-bound RNA

Unbound samples were collected for RNA extractions, three volumes of peqGOLD TriFast™ FL reagent were added, and samples were stored at −80 °C until further processing. The beads were washed four times with wash buffer and proteins with bound RNAs were released by incubation with 5 μL recombinant TEV protease (1 mg/mL) in 250 μL of wash buffer for 90 min at 20 °C. For collecting the elution fractions, the beads were concentrated on one side of the tube, the supernatant was transferred to a fresh tube, three volumes of peqGOLD TriFast™ FL reagent were added, and samples were stored at −80 °C until further processing.

### RNA isolation

Samples frozen at −80 °C in TriFast reagent were thawed and incubated for 5 min at room temperature to ensure complete dissociation of ribonuclear complexes. Afterwards, 200 μL of chloroform were added per sample, and the tubes were shaken vigorously by hand for 15 s. Subsequent to 3 min of incubation at room temperature, samples were centrifuged at 12,000 rpm for 10 min at 4 °C. Following centrifugation, the aqueous phase was transferred to a fresh tube, and 7 μL of glycogen solution (10 mg/mL) were added, as well as 500 μL of isopropanol. The samples were mixed briefly by vortexing and incubated at room temperature for 10 min, before being subjected to another centrifugation step at 12,000 rpm for 15 min at 4 °C. The pellet was subjected to two sequential washing steps with 70% and absolute ethanol, respectively. Afterwards, the supernatant was removed and the pellet was dried at room temperature for 10 min. Purified RNA was then dissolved in nuclease-free water by incubation at 50 °C for 5 min, and the RNA concentration was determined spectrophotometrically.

### Analysis of circular RNAs

For this purpose, 1 × 10^8^ procyclic forms were harvested and total RNA was extracted according to the standard protocol. Afterwards, 30 μg of RNA were treated with 20 U of DNase I in 200 μL in presence of RNaseIN for 30 min at 25 °C. The reaction was terminated by phenol-chloroform extraction and ethanol precipitation of the RNA. The pellet was dissolved in 32 μL of RNase-free water and incubated with 100 pmol of an oligonucleotide complementary to the first 15 nt of the spliced leader (5’ TCTAATAATAGCGTT 3’) at 37 °C for 5 min. RNase H buffer was added to reach a concentration of 1×, along with 5 U of RNase H, and the reaction was then incubated for 1 h at 37 °C. The RNA was then purified and precipitated with ethanol according to the standard protocol. Subsequently, 10 μg of RNA were circularised by incubation at 16 °C for 16 h in a reaction volume of 400 μL containing 40 U of T4 RNA and 80 U of RNaseIN. For reverse transcription, ~2 μg of RNA were incubated with 50 pmol of GPEET-specific reverse primer, 200 U SuperScript III RT (Invitrogen), and 40 U RNase Inhibitor in a 20 μL reaction volume.

PCR amplification was performed in a 50 μL reaction using 1 μL of cDNA (5 %), 10 pmol each of forward and reverse primers, 2.5 U Taq DNA polymerase (30 s at 95 °C, 30 s at 52 °C, and 45 s at 72 °C). The PCR products were gel-purified and cloned into EamI-digested p2T7 vector. Clones were subjected to bluewhite selection and analysed by sequencing.

### Expression analysis by RT-qPCR

For this, cDNA was synthesized from unbound and bound fractions of RNA-immunoprecipitations using the Maxima First Strand cDNA Synthesis Kit (Thermo Scientific) according to the manufacturer’s instructions. Afterwards, expression of the mRNAs of interest was analysed in triplicates by qPCR using the Luna^®^ Universal qPCR Master Mix and specific primers (Supplementary table S3). Data were analysed by the 2^-ΔΔCt^ method and plotted with GraphPad Prism software (version 6).

### RNA sequencing and data analysis

RNA-seq was done at the CellNetworks Deep Sequencing Core Facility at the University of Heidelberg. For library preparation, NEBNext Ultra RNA Library Prep Kit for Illumina (New England BioLabs Inc.) was used. The libraries were multiplexed (6 samples per lane) and sequenced with a Nextseq 550 system, generating 75 bp single-end sequencing reads. Dta are deposited as E-MTAB-10453

The quality of the data was checked using FastQC. After primer removal using Cutadapt, the data were aligned to the TREU 927 reference genome using custom scripts, as described previously (Mugo and Clayton 2017; Melo do Nascimento et al. 2021).

### *In silico* identification of uridylation targets

Identification of potentially uridylated transcripts was performed RNA sequencing data from four sets of bloodstream form and procyclic form data each (E-MTAB-3793; E-GEOD-79208; E-MTAB-8062; E-MTAB-7996; E-MTAB-3335). The sequence reads, which were 75 nt on average, were analysed for the presence of at least 15 As followed by > 5 Us. Afterwards, the 3’ UTRs preceding the 3’ uridylated transcripts served to identify the corresponding transcript through BLAST search in the TriTryp database.

### Digitonin fractionation

10^8^ cells were collected by centrifugation (950 × *g*, 8 min) and washed three times in trypanosome homogenization buffer, THB (25 mM Tris-Cl [pH 7.8], 1 mM EDTA, 150 mM NaCl, 0.3 M sucrose). The washes consisted of centrifugation (950 × *g*, 3 min, 4 °C) and resuspension by gentle agitation of the tube instead of pipetting. The cells were eventually resuspended in 100 μL of THB containing 2 μg/mL leupeptin, 1 mM DTT and always placed on ice. Using a stock of 0.9 mg/mL of digitonin (dissolved in THB 95 °C, 5 min), aliquots of different digitonin concentrations were prepared and their volumes adjusted to be similar to each other and set at room temperature. 10^7^ cells (10 μL) were pipetted directly into the digitonin aliquots (at most, three tubes at a time) and incubated for 4 min at 25 °C. Subsequently, the tubes were centrifuged (15,871 × *g*, 1.5 min, 4 °C) and the supernatants and pellets separated and dissolved in laemmli buffer (125 mM Tris-HCl pH 6.8, 4 % SDS, 15 mM EDTA, 10 % β-mercaptoethanol, 20 % glycerol, 0.1 % bromophenol blue). The samples were separated on a 10 % SDS-PAGE gel and analysed by western blotting (see above).

### Yeast-2-hybrid assays

For testing direct protein-protein interactions, the Matchmaker Yeast Two-Hybrid System (Clontech) was used according to the manufacturer’s instructions. To that end, ORFs of interest (TUT3, 4EIP, and Dis3L2) were PCR-amplified from genomic DNA and cloned into both pGBKT7 and pGADT7 plasmids. Prey and bait plasmids were co-transformed pairwise into AH109 yeast strains, and selected initially on double drop-out (DDO) plates (i.e., SD medium lacking Trp and Leu) or quadruple drop-out (QDO) plates (i.e., lacking Trp, Leu, His and Ade). Growth on QDO plates indicated positive interactions. The interaction between p53 and SV40 large T-antigen and the combination of LaminC and SV40 large T-antigen served as positive and negative controls, respectively. Western blotting was used to confirm expression of c-myc-tagged BD-domain proteins and HA-tagged AD-domain proteins.

### Protein detection by western blotting

For western blotting, 1-5 × 10^6^ cells were lysed in 1× Laemmli buffer, separated by 8-12 % SDS-PAGE, and processed as described elsewhere (Klein et al. 2015). The following antibodies were used for specific protein detection: anti-4E1 serum (1:2,000, rabbit; kind gift from Osvaldo de Melo Neto), anti-S9 serum (1:20,000, rat; loading control), anti-myc (1:5,000, mouse; Y2H), anti-p-GPEET (1:2,000, mouse, Cedarlane; western blotting), anti-V5 (1:2,000, mouse, Biorad; western blotting), anti-PAP (1:20,000, rabbit, Sigma; western blotting), anti-aldolase serum (1:50,000, rabbit; western blotting), anti-EP (1:2,000, mouse, Cedarlane; western blotting), and anti-CAT serum (1:2,000, rabbit), anti-rabbit IgG (pull-downs).

## ACKNOWLEDGEMENTS

We thank Ute Leibfried and Claudia Helbig for technical assistance, Georg Stoecklin for useful discussions, and Osvaldo de Melo Neto (Fiocruz, Recife, Brazil) for providing plasmids and antibodies. Sequencing was done by David Ibberson (Deep Sequencing Core Facility), and Thomas Ruppert and Sabine Merker (Core Facility for Mass Spectrometry) performed the mass spectrometry.

KM was initially supported by a scholarship from the Deutscher Akademischer Austauschdienst. This project was mainly supported by grant number Cl112/30 from the Deutsche Forschungsgemeinschaft.

## AUTHOR CONTRIBUTIONS

KKM contributed data on TUT3 localization, KO and RNAi studies (Fig. 4, Supplementary Fig. S2 A and B, S3, and S4), as well as the corresponding text passages. FF performed all other experiments. CC was responsible for conceptualisation, funding acquisition, supervision, and project administration. CC and FF analysed the data and wrote the manuscript.

## Supplementary Figures

**Fig. S1.**
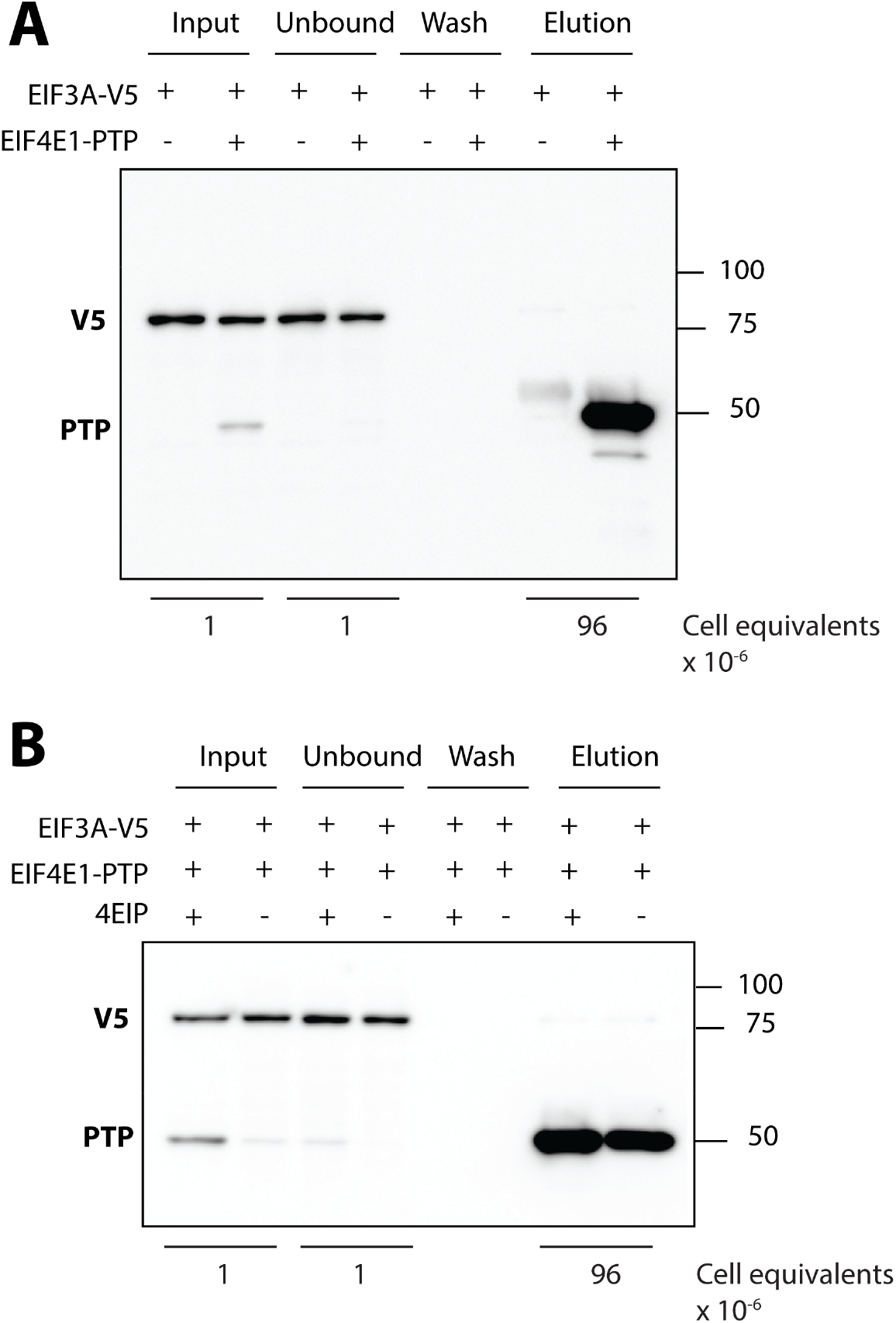
EIF4E1 does not pull down EIF3A in absence and presence of 4EIP. **(A)** PTP-tagged EIF4E1 was pulled down from 1 × 10^8^ bloodstream forms (BSFs) of *Trypanosoma brucei*. Enrichment of V5-tagged EIF3A in the different fractions was analysed by western blotting. **(B)** Same experiment as described in **(A)**, including cells lacking 4EIP.

**Fig. S2.**
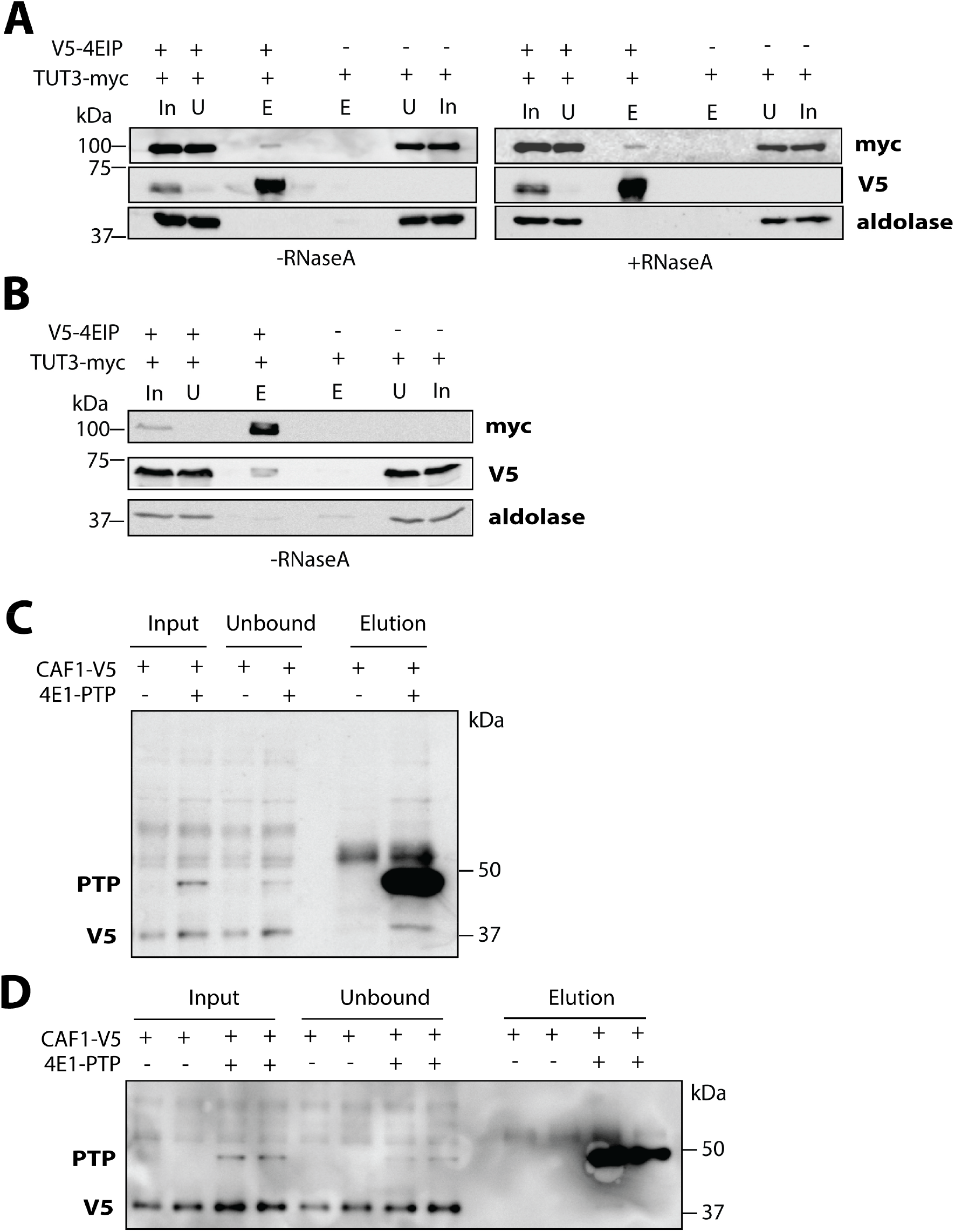
Recruitment of TUT3 and CAF1 to EIF4E1/4EIP complexes. **(A)** Pull-downs of V5-tagged 4EIP from bloodstream form (BSF) *Trypanosoma brucei* parasites in absence or presence of RNase A (left and right panels, respectively), followed by detection of TUT3-myc enrichment by western blotting. **(B)** Pull-down of myc-tagged TUT3 from bloodstream form *T. brucei* parasites, followed by detection of 4EIP-V5 by western blotting. **(C, D)** Two representative replicates of pull-downs of PTP-tagged EIF4E1 from bloodstream form *T. brucei* parasites, followed by detection of CAF1-V5 by western blotting.

**Fig. S3.**
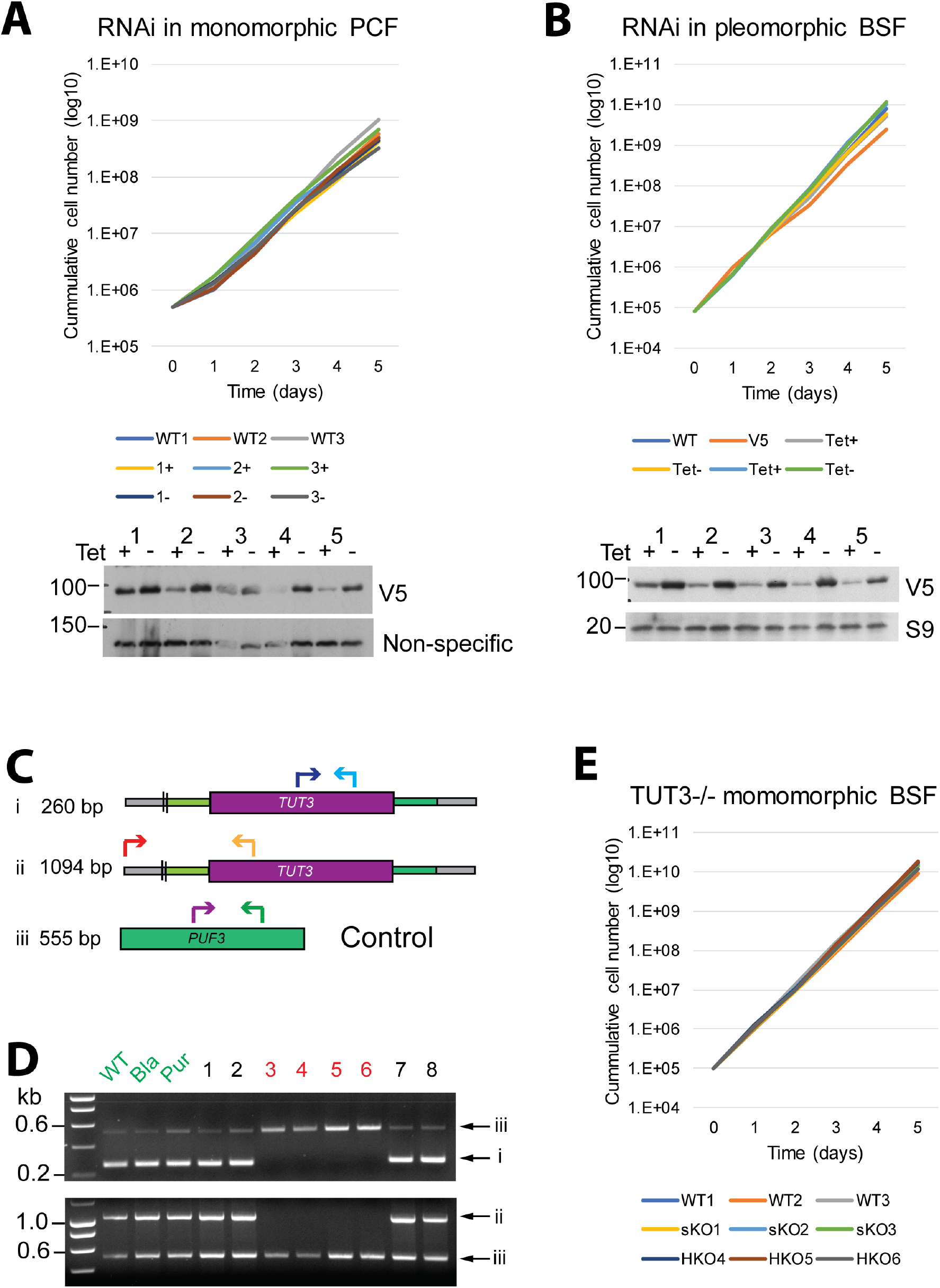
Effects of TUT3 depletion or elimination. **(A)** Knockdown of TUT3 was induced in monomorphic procyclic forms (PCFs) cultured in presence of tetracycline (+), with wild-type (WT) cells and RNAi clones cultured in absence of tetracycline serving as controls. The cell numbers were determined over the course of 5 days (upper panel), and knockdown efficiency was determined by western blotting (lower panel). **(B)** Knockdown of TUT3 was induced in bloodstream forms (BSFs) cultured in presence of tetracycline (+), with wild-type (WT) cells and RNAi clones cultured in absence of tetracycline serving as controls. The cell numbers were determined over the course of 5 days (upper panel), and knockdown efficiency was determined by western blotting (lower panel). **(C)** Primer pairs used for amplifying a 260 bp fragment within the TUT3 ORF (i) or a fragment spanning the 5’-UTR and the ORF (ii). Primers for amplification of the PUF3 gene were used as a control (iii). **(D)** Diagnostic polymerase chain reaction for the *TUT3* gene. Each PCR contained primers to amplify *PUF3* (purple, green, band iii) as a positive control. Product (i) was obtained from a primer pair (blue, cyan) that hybridised within the *TUT3* open reading frame. Product (ii) was obtained using one primer that targets the ORF (orange) and another (red) that binds upstream of *TUT3* to a sequence absent from the knockout plasmid. **(E)** The growth of monomorphic bloodstream forms with knockout of either a single copy or both copies of the TUT3 coding sequence (SKO and HKO, respectively) was monitored over the course of 5 days.

**Fig. S4.**
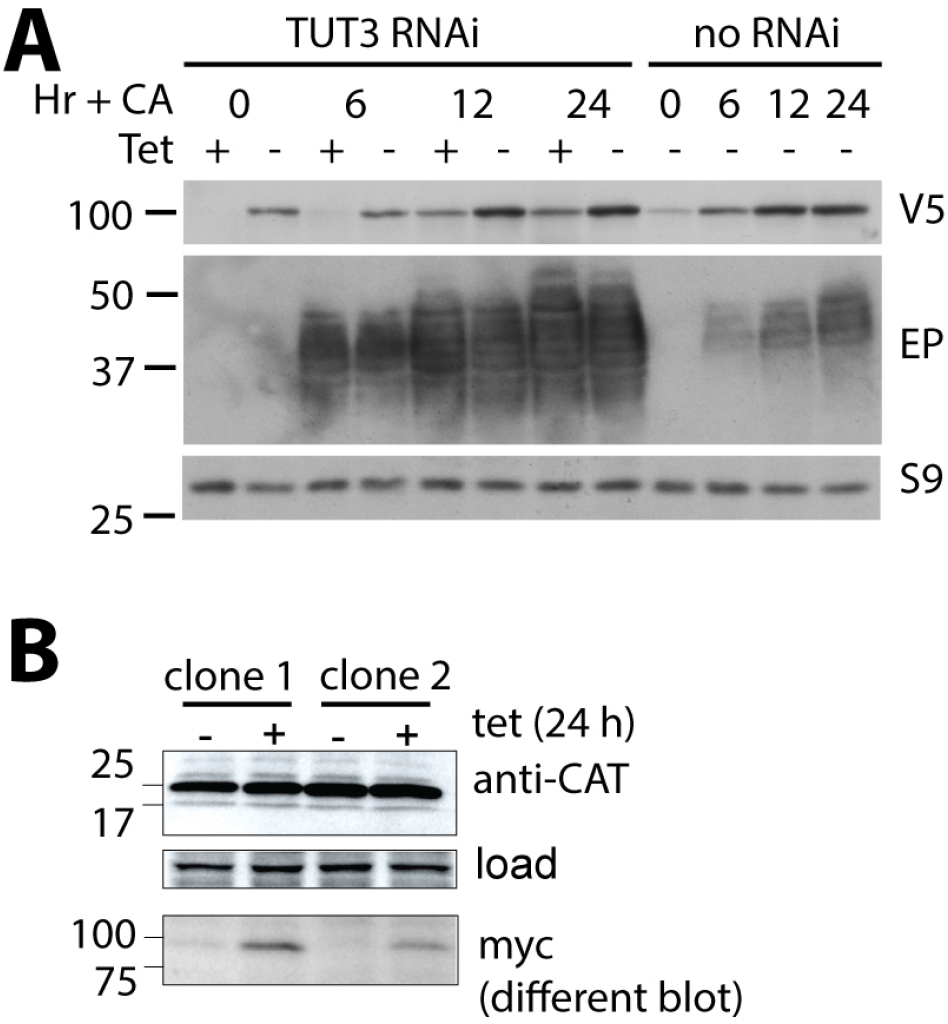
Differentiation of TUT3-depleted cells and effect of tethering. **(A)** Differentiation-competent T. brucei (EATRO1125 strain) with one V5-tagged TUT3 gene and RNAi targeting *TUT3* were grown to a maximum density of 2 × 10^6^ cells/mL. 6 mM *cis*-aconitate (CA) was then added, and the temperature was reduced to 27 °C to induce differentiation. Cultures were grown with or without tetracycline, with the drug being included for 24 h before addition of *cis*-aconitate. For each time point, 5 × 10^6^ cells were collected for western blot measurement of V5-TUT3, Ep procyclin, and a control rpotein, ribosomal protein S9. **(B)** Tethering of lambdaN-TUT3-myc has no effect on a box-B-containing reporter mRNA encoding chloramphenicaol acetyltransferase (CAT). Expression of lambdaN-TUT3-myc was induced for 24h then CATwas measured by Western blotting. A protein that cross-reacts with the anti-CAT antibody served as a loading control.

**Fig. S5.**
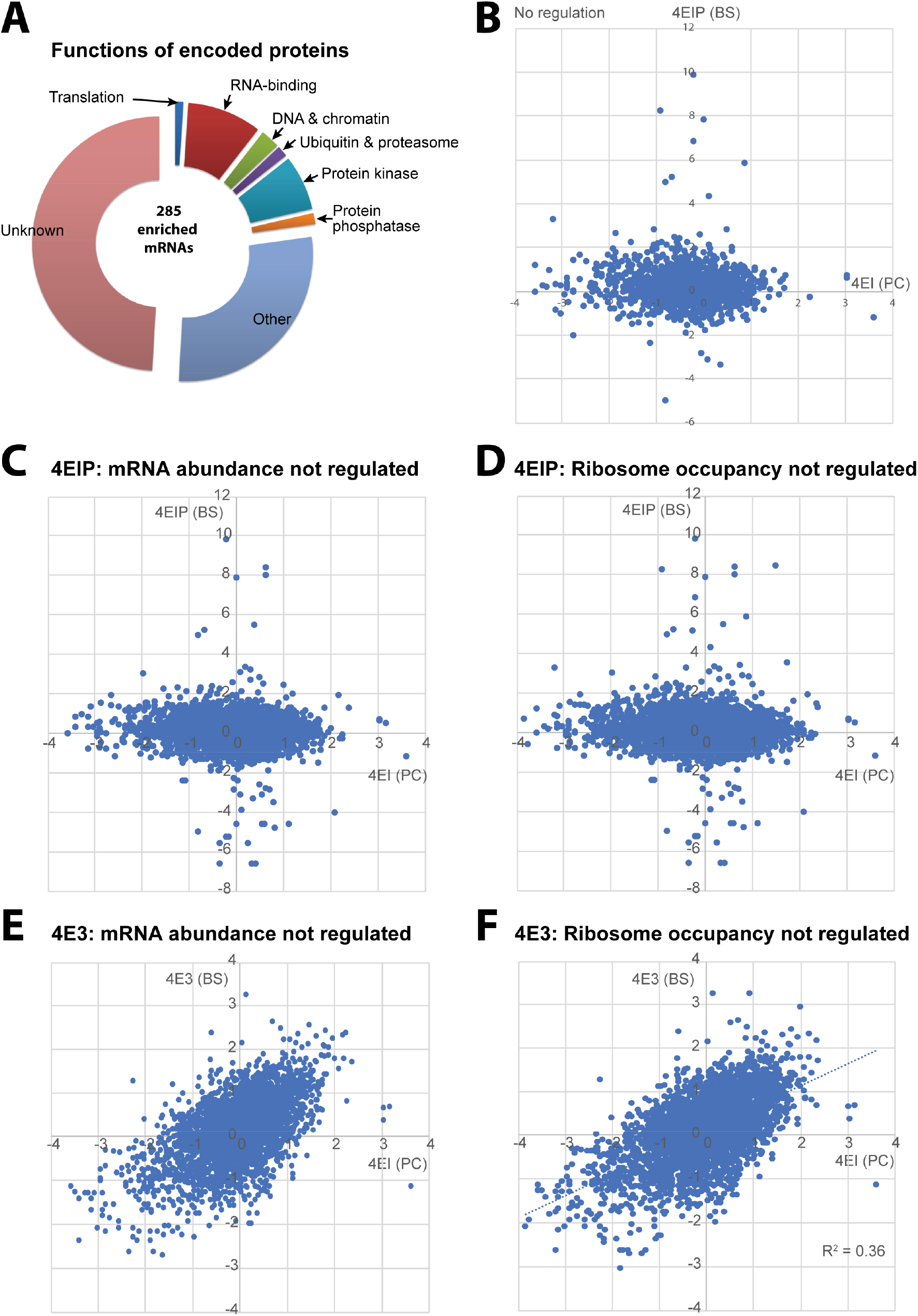
EIF4E1-associated mRNAs. **(A)** Functions of proteins encoded by the 285 mRNAs that were most enriched in the EIF4E1 pull-downs. **(B)** The mRNAs bound to EIF4E1 in procyclic forms were compared with those bound to 4EIP in bloodstream forms. The analysis was restricted to mRNAs that show less than 1.5-fold differences between bloodstream and procyclic forms in both mRNA abundance and ribosome occupancy (Antwi et al. 2016). **(C)** As (B) but including only mRNAs that show less than 1.5-fold differences between bloodstream and procyclic forms in mRNA abundance. **(D)** As (B) but including only mRNAs that show less than 1.5-fold differences between bloodstream and procyclic forms in ribosome occupancy. **(E)** The mRNAs bound to EIF4E1 in procyclic forms were compared with those bound to EIF4E3 in bloodstream forms. The analysis includes only mRNAs that show less than 1.5-fold differences between bloodstream and procyclic forms in mRNA abundance. **(F)** The mRNAs bound to EIF4E1 in procyclic forms were compared with those bound to EIF4E3 in bloodstream forms. The analysis includes only mRNAs that show less than 1.5-fold differences between bloodstream and procyclic forms in ribosome occupancy.

## Supplementary file

**File S6 Sequencing of poly(A)-tailed GPEET mRNAs**

*GPEET* mRNA 3’-ends in normal and glucose-adapted WT procyclic forms (PCFs) were examined by cap removal, circularization, selective amplification, and DNA sequencing. Identified poly(A)-tailed *GPEET* mRNAs from low- and high-glucose-grown cells are shown; 20 sequences were analysed each.

## Supplementary Tables

**Table S1 Proteins associated with EIF4E1 in absence and presence of 4EIP**

A detailed legend is on the first sheet of the Table.

**Table S2 mRNAs associated with EIF4E1 in procyclic forms, in the absence and presence of 4EIP**

A detailed legend is on the first sheet of the Table.

**Table S3 Plasmids and oligonucleotides**

## REFERENCES

Alsford S, Turner D, Obado S, Sanchez-Flores A, Glover L, Berriman M, Hertz-Fowler C, Horn D. 2011. High throughput phenotyping using parallel sequencing of RNA interference targets in the African trypanosome. Genome Res 21: 915–924.

Amaya Ramirez CC, Hubbe P, Mandel N, Bethune J. 2018. 4EHP-independent repression of endogenous mRNAs by the RNA-binding protein GIGYF2. Nucleic Acids Res 46: 5792–5808.

Antwi EB, Haanstra JR, Ramasamy G, Jensen B, Droll D, Rojas F, Minia I, Terrao M, Merce C, Matthews K et al. 2016. Integrative analysis of the Trypanosoma brucei gene expression cascade predicts differential regulation of mRNA processing and unusual control of ribosomal protein expression. BMC Genomics 17: 306.

Aphasizhev R, Aphasizheva I. 2011. Uridine insertion/deletion editing in trypanosomes: a playground for RNA-guided information transfer. WIRES RNA 2: 669–685.

Aphasizhev R, Aphasizheva I, Simpson L. 2004. Multiple terminal uridylyltransferases of trypanosomes. FEBS Lett 572: 15–18.

Barbour AG, Restrepo BI. 2000. Antigenic variation in vector-borne pathogens. Emerg Infect Dis 6: 449–457.

Baron N, Tupperwar N, Dahan I, Hadad U, Davidov G, Zarivach R, Shapira M. 2021. Distinct features of the Leishmania cap-binding protein LeishIF4E2 revealed by CRISPR-Cas9 mediated hemizygous deletion. PLoS Negl Trop Dis 15: e0008352.

Brecht M, Parsons M. 1998. Changes in polysome profiles accompany trypanosome development. Mol Biochem Parasitol 97: 189–198.

Bringaud F, Riviere L, Coustou V. 2006. Energy metabolism of trypanosomatids: adaptation to available carbon sources. Mol Biochem Parasitol 149: 1–9.

Cho PF, Poulin F, Cho-Park YA, Cho-Park IB, Chicoine JD, Lasko P, Sonenberg N. 2005. A new paradigm for translational control: inhibition via 5’-3’ mRNA tethering by Bicoid and the eIF4E cognate 4EHP. Cell 121: 411–423.

Clayton CE. 2016. Gene expression in Kinetoplastids. Curr Opin Microbiol 32: 46–51.

Dean S, Marchetti R, Kirk K, Matthews KR. 2009. A surface transporter family conveys the trypanosome differentiation signal. Nature 459: 213–217.

Dean S, Sunter JD, Wheeler RJ. 2017. TrypTag.org: A Trypanosome Genome-wide Protein Localisation Resource. Trends Parasitol 33: 80–82.

Dejung M, Subota I, Bucerius F, Dindar G, Freiwald A, Engstler M, Boshart M, Butter F, Janzen CJ. 2016. Quantitative proteomics uncovers novel factors involved in developmental dfferentiation of *Trypanosoma brucei*. PLoS pathogens 12: e1005439.

Erben E, Leiss K, Liu B, Gil DI, Helbig C, Clayton C. 2021. Insights into the functions and RNA binding of Trypanosoma brucei ZC3H22, RBP9 and DRBD7. Parasitology: 1–10.

Erben ED, Fadda A, Lueong S, Hoheisel JD, Clayton C. 2014. A genome-wide tethering screen reveals novel potential post-transcriptional regulators in Trypanosoma brucei. PLoS Pathog 10: e1004178.

Fadda A, Ryten M, Droll D, Rojas F, Farber V, Haanstra JR, Merce C, Bakker BM, Matthews K, Clayton C. 2014. Transcriptome-wide analysis of trypanosome mRNA decay reveals complex degradation kinetics and suggests a role for co-transcriptional degradation in determining mRNA levels. Mol Microbiol 94: 307–326.

Firczuk H, Kannambath S, Pahle J, Claydon A, Beynon R, Duncan J, Westerhoff H, Mendes P, McCarthy JE. 2013. An in vivo control map for the eukaryotic mRNA translation machinery. Mol Syst Biol 9: 635.

Freire ER, Moura DMN, Bezerra MJR, Xavier CC, Morais-Sobral MC, Vashisht AA, Rezende AM, Wohlschlegel JA, Sturm NR, de Melo Neto OP et al. 2018. Trypanosoma brucei EIF4E2 cap-binding protein binds a homolog of the histone-mRNA stem-loop-binding protein. Curr Genet 64: 821–839.

Freire ER, Sturm NR, Campbell DA, de Melo Neto OP. 2017. The Role of Cytoplasmic mRNA Cap-Binding Protein Complexes in Trypanosoma brucei and Other Trypanosomatids. Pathogens 6.

Fritz M, Vanselow J, Sauer N, Lamer S, Goos C, Siegel T, Subota I, Schlosser A, Carrington M, Kramer S. 2015. Novel insights into RNP granules by employing the trypanosome’s microtubule skeleton as a molecular sieve. Nucleic Acids Res 43: 8013–8032.

Fu R, Olsen MT, Webb K, Bennett EJ, Lykke-Andersen J. 2016. Recruitment of the 4EHP-GYF2 cap-binding complex to tetraproline motifs of tristetraprolin promotes repression and degradation of mRNAs with AU-rich elements. RNA 22: 373–382.

Gunasekera K, Wuthrich D, Braga-Lagache S, Heller M, Ochsenreiter T. 2012. Proteome remodelling during development from blood to insect-form Trypanosoma brucei quantified by SILAC and mass spectrometry. BMC genomics 13: 556.

Hernandez G, Altmann M, Sierra JM, Urlaub H, Diez del Corral R, Schwartz P, Rivera-Pomar R. 2005. Functional analysis of seven genes encoding eight translation initiation factor 4E (eIF4E) isoforms in Drosophila. Mech Dev 122: 529–543.

Hernandez G, Proud CG, Preiss T, Parsyan A. 2012. On the Diversification of the Translation Apparatus across Eukaryotes. Comp Funct Genomics 2012: 256848.

Hickey KL, Dickson K, Cogan JZ, Replogle JM, Schoof M, D’Orazio KN, Sinha NK, Hussmann JA, Jost M, Frost A et al. 2020. GIGYF2 and 4EHP Inhibit Translation Initiation of Defective Messenger RNAs to Assist Ribosome-Associated Quality Control. Mol Cell 79: 950–962 e956.

Horn D. 2014. Antigenic variation in African trypanosomes. Mol Biochem Parasitol 195: 123–129.

Igreja C, Izaurralde E. 2011. CUP promotes deadenylation and inhibits decapping of mRNA targets. Genes Dev 25: 1955–1967.

Jensen BC, Ramasamy G, Vasconcelos EJ, Ingolia NT, Myler PJ, Parsons M. 2014. Extensive stage-regulation of translation revealed by ribosome profiling of *Trypanosoma brucei*. BMC genomics 15: 911.

Juszkiewicz S, Slodkowicz G, Lin Z, Freire-Pritchett P, Peak-Chew SY, Hegde RS. 2020. Ribosome collisions trigger cis-acting feedback inhibition of translation initiation. Elife 9.

Kamenska A, Lu WT, Kubacka D, Broomhead H, Minshall N, Bushell M, Standart N. 2014a. Human 4E-T represses translation of bound mRNAs and enhances microRNA-mediated silencing. Nucleic Acids Res 42: 3298–3313.

Kamenska A, Simpson C, Standart N. 2014b. eIF4E-binding proteins: new factors, new locations, new roles. Biochem Soc Trans 42: 1238–1245.

Klein C, Terrao M, Inchaustegui Gil D, Clayton C. 2015. Polysomes of *Trypanosoma brucei:* association with initiation factors and RNA-binding proteins. PloS ONE 10: e0135973.

Knusel S, Roditi I. 2013. Insights into the regulation of GPEET procyclin during differentiation from early to late procyclic forms of Trypanosoma brucei. Mol Biochem Parasitol 191: 66–74.

Kropiwnicka A, Kuchta K, Lukaszewicz M, Kowalska J, Jemielity J, Ginalski K, Darzynkiewicz E, Zuberek J. 2015. Five eIF4E isoforms from Arabidopsis thaliana are characterized by distinct features of cap analogs binding. Biochem Biophys Res Commun 456: 47–52.

Lackey PE, Welch JD, Marzluff WF. 2016. TUT7 catalyzes the uridylation of the 3’ end for rapid degradation of histone mRNA. RNA 22: 1673–1688.

Lueong S, Merce C, Fischer B, Hoheisel JD, Erben ED. 2016. Gene expression regulatory networks in Trypanosoma brucei: insights into the role of the mRNA-binding proteome. Mol Microbiol 100: 457–471.

Mabille D, Cardoso Santos C, Hendrickx R, Claes M, Takac P, Clayton C, Hendrickx S, Hulpia F, Maes L, Van Calenbergh S et al. 2021. 4E Interacting Protein as a Potential Novel Drug Target for Nucleoside Analogues in Trypanosoma brucei. Microorganisms 9.

Mader S, Lee H, Pause A, Sonenberg N. 1995. The translation initiation factor eIF-4E binds to a common motif shared by the translation factor eIF-4 gamma and the translational repressors 4E-binding proteins. Mol Cell Biol 15: 4990–4997.

Mantilla BS, Marchese L, Casas-Sanchez A, Dyer NA, Ejeh N, Biran M, Bringaud F, Lehane MJ, Acosta-Serrano A, Silber AM. 2017. Proline Metabolism is Essential for Trypanosoma brucei brucei Survival in the Tsetse Vector. PLoS Pathog 13: e1006158.

Meleppattu S, Arthanari H, Zinoviev A, Boeszoermenyi A, Wagner G, Shapira M, Leger-Abraham M. 2018. Structural basis for LeishIF4E-1 modulation by an interacting protein in the human parasite Leishmania major. Nucleic Acids Res 46: 3791–3801.

Meleppattu S, Kamus-Elimeleh D, Zinoviev A, Cohen-Mor S, Orr I, Shapira M. 2015. The eIF3 complex of Leishmania-subunit composition and mode of recruitment to different cap-binding complexes. Nucleic Acids Res 43: 6222–6235.

Melo do Nascimento L, Egler F, Arnold K, Papavasiliou N, Clayton C, Erben E. 2021. Functional insights from a surface antigen mRNA-bound proteome. Elife 10.

Morita M, Ler LW, Fabian MR, Siddiqui N, Mullin M, Henderson VC, Alain T, Fonseca BD, Karashchuk G, Bennett CF et al. 2012. A novel 4EHP-GIGYF2 translational repressor complex is essential for mammalian development. Mol Cell Biol 32: 3585–3593.

Mugo E, Clayton C. 2017. Expression of the RNA-binding protein RBP10 promotes the bloodstream-form differentiation state in Trypanosoma brucei. PLoS Pathog 13: e1006560.

Pays E, Hanocq-Quertier J, Hanocq F, Van Assel S, Nolan D, Rolin S. 1993. Abrupt RNA changes precede the first cell division during the differentiation of Trypanosoma brucei bloodstream forms into procyclic forms in vitro. Mol Biochem Parasitol 61: 107–114.

Pestova TV, Kolupaeva VG, Lomakin IB, Pilipenko EV, Shatsky IN, Agol VI, Hellen CU. 2001. Molecular mechanisms of translation initiation in eukaryotes. Proc Natl Acad Sci U S A 98: 7029–7036.

Peter D, Weber R, Sandmeir F, Wohlbold L, Helms S, Bawankar P, Valkov E, Igreja C, Izaurralde E. 2017. GIGYF1/2 proteins use auxiliary sequences to selectively bind to 4EHP and repress target mRNA expression. Genes Dev 31: 1147–1161.

Rom E, Kim HC, Gingras AC, Marcotrigiano J, Favre D, Olsen H, Burley SK, Sonenberg N. 1998. Cloning and characterization of 4EHP, a novel mammalian eIF4E-related cap-binding protein. J Biol Chem 273: 13104–13109.

Ruscica V, Bawankar P, Peter D, Helms S, Igreja C, Izaurralde E. 2019. Direct role for the Drosophila GIGYF protein in 4EHP-mediated mRNA repression. Nucleic Acids Res 47: 7035–7048.

Shiber A, Doring K, Friedrich U, Klann K, Merker D, Zedan M, Tippmann F, Kramer G, Bukau B. 2018. Cotranslational assembly of protein complexes in eukaryotes revealed by ribosome profiling. Nature 561: 268–272.

Terrao M, Marucha KK, Mugo E, Droll D, Minia I, Egler F, Braun J, Clayton C. 2018. The suppressive cap-binding complex factor 4EIP is required for normal differentiation. Nucleic Acids Res 46: 8993–9010.

Trippe R, Guschina E, Hossbach M, Urlaub H, Luhrmann R, Benecke BJ. 2006. Identification, cloning, and functional analysis of the human U6 snRNA-specific terminal uridylyl transferase. RNA 12: 1494–1504.

Tupperwar N, Meleppattu S, Shrivastava R, Baron N, Gilad A, Wagner G, Leger-Abraham M, Shapira M. 2020. A newly identified Leishmania IF4E-interacting protein, Leish4E-IP2, modulates the activity of cap-binding protein paralogs. Nucleic Acids Res 48: 4405–4417.

Tupperwar N, Shrivastava R, Shapira M. 2019. LeishIF4E1 Deletion Affects the Promastigote Proteome, Morphology, and Infectivity. mSphere 4.

Vassella E, Probst M, Schneider A, Studer E, Renggli CK, Roditi I. 2004. Expression of a major surface protein of Trypanosoma brucei insect forms is controlled by the activity of mitochondrial enzymes. Mol Biol Cell 15: 3986–3993.

Vickerman K. 1978. Antigenic variation in trypanosomes. Nature 273: 613–617.

Viegas SC, Silva IJ, Apura P, Matos RG, Arraiano CM. 2015. Surprises in the 3’-end: ‘U’ can decide too! Febs j 282: 3489–3499.

Weber R, Chung MY, Keskeny C, Zinnall U, Landthaler M, Valkov E, Izaurralde E, Igreja C. 2020. 4EHP and GIGYF1/2 Mediate Translation-Coupled Messenger RNA Decay. Cell Rep 33: 108262.

Zinoviev A, Leger M, Wagner G, Shapira M. 2011. A novel 4E-interacting protein in Leishmania is involved in stage-specific translation pathways. Nucleic Acids Res 39: 8404–8415.

