## Supplemental file S6 for "Mechanisms of translation repression by the EIF4E1-4EIP cap-binding complex of *Trypanosoma brucei*: potential roles of the NOT complex and a terminal uridylyl transferase"

### Supplementary file S6

**Start of 5' UTR**

**Part of spliced leader**

**polyA tail**

**End of 3' UTR**

**20 analyzed each**

### Low glucose:

CGCCTCGGAGGAACGAAACCCCTTTGAAAAGGTGGCTTTCGTTTATATATTCTCCATCTGGTGCATCGTGTTTATTTCGTATTATTCCCTGTCAA
AAGCTGCGGATATTCATTTAATATTCTTTTCGTTATATATTTTTAGTTCTATCCTTTATTTTATCCGTTTTAATCGTCTTCTGAGACCCACAGCCCT
GTAGATTTCTGTGATGTTTCGGTTGCGTATTCCATAATTTTAANCGTTCCACTCCTATTTTTTTCATTCCTTTGAATTTGGATCTTNANNAAAATA
AAAAAAAAAAAAAAAAAAAAAAAAAAAAAAAAAGAAAAAAAAAAAAAAAAAAAAAAAAGTTANAACAGTTTCTGTACCATATNGACTTCAN
TTACNCCAAAAAGNAAAATTCNCAAGGGCNCCTCGTTCCCTTTNTCNGCNCGCTATTCTTCTGTTCAGCGCGAACNAAGGTCGTTAACNCGNG
GGGGGGCCCGGGGANAT

ATAGCGAGCAGATAAAGGGAACGAGGTGCCATTGTGAATTTTACTTTTTGGTGTAATTGAAGTCAATATAGTACAGAAACTGTTCTAATAATTT
TTTTTTTTTTTTTTTTTTTTTTTTTTTTTTTTTTTTTTTTTTTTTTTTTTTTTTTTTTTTTTTTTTTTTTTNAAA

GTGTAAAGCGCCTCGGAGGAACGAAACCCCTTTGAAAAGGTGGCTTTCGTTTATATATTCTCCATCTGGTGCATCGTGTTTATTTCGTATTATTC
CCTGTCAAAAGCTGCGGATATTCATTTAATATTCTTTTCGTTATATATTTTTAGTTCTATCCTTTATTTTATCCGTTTTAATCGTCTTCTGAGACCCA
CAGCCCTGTAGATTTCTGTGATGTTTCGGTTGCGTATTCCATAATTTTAAGCGTTTCACTTCTATTTTTTTTCATTCCTTTGAATTTGGATCTTAAA
AAAAAAAAAAAAAAAAAAAAAAAAAAAAAAAAAAAAAAAAAAAAAAAAAAAAAAAAAAAAAAAAAAAAAAAAAAAAAAAAAAANTT

CGGATGCAAGCGTGTAAAGCGCCTCGGAGGAACGAAACCCTTTGAAAAGGTTCCTTTCATTTATATCGCCTCCATATGGTGCATCGTGTTTGTT
TCCTGCTGTTTCTTGTAAAACAAGTGTGGACATTCATTTAATATTTTTTCGTTATATTTTTTTGGTGACATCCTTTCTAATGCCTTATTGACCATC
GCCTGAGACCCACAGCCCTGTAGATTTCTGTGATGTTTCGGTTGCGTATTCCATAATTTTAANCGTTTCACTTCTATTTTTTTCATTCCTTTGAATT
TGGATCTTAAAAAAAAAAAAAAAAAAAAAAAAAAAAAAAAAAAAAAAAAAAAAAAAAAAAAAAAAAAAAAAAAAAAAAAAAAAAAAAAAA
AAAAAAAAAAAAAAAAAAAAAAAAAAAAAAAA

TAGCGAGCAGATAAAGGGAACGAGGTGCCATTGTGAATTTTACTTTTTGGTGTAATTGAAGTCAATATAGTACAGAAACTGTTCTAATTTTTTT
TTTTTTTTTTTTTTTTTTTTTTTTTTTTTTTTTTTTTTTTTTTTTTTTTTTTTTTTTTTTTTTTTTTTTTTTTTTTTTTTTTTTTTTTTTTTTTTTTTTTTTTTT
TTTTTTTTTTTTTTTN

GCGNGTAAAGCGCCTCGGAGGAACGAAACCCTTTGAAAAGGTTCCTTTCATTTATATCGCCTCCATATGGTGCATCGTGTTTGTTTCCTGCTGT
TTCTTGTAAAACAAGTGTGGACATTCATTTAATATTTTTTCGTTATATTTTTTTGGTGACATCCTTTCTAATGCCTTATTAACCATCGCCTGAGACC

CACAGCCCTGTAGATTTCTGTGATGTTTCGGTTGCGTATTCCATAATTTTAAGCGTTTCACTTCTATTTTTTTTCATTCCTTTGAATTTGGATCTTA
AAAAAAAAAAAAAAAAAAAAAAAAAAAAAAAAAAAAAAAAAAAAAAAAAAAAAAAAAAAAAAAAAAAAAAANG

ATNNCGAGCAGATAAAGGGAACGAGGTGCCATTGTGAATTTTACTTTTTGGTGTAATTGAAGTCAATATAGTACAGAAACTGTTCTAATATTTT
TTTTTTTTTTTTTTTTTTTTTTTTTTTTTTTTTTTTTTTTTTTTTTTTTTTTTTTTTTTTTTTTTTTTTTTTTTTTTTTTTTTTTTTTTNNA

AAAGGGAACNAGGNGCCATTGTGAATTTTACTTTTTGGTGTAATTGAAGTCAATATAGTACAGAAACTGTTCTAATACTTTTTTTATTTTTTTTT
TTTTTTTTTTTTTTTTTTTTTTTTTTTTTTTTNCNTTTNCNTTTTTTTTTTTTTTTTTTTTTTTTTT

**High glucose:**

GCGGATGCAAGCGTGTAAAGCGCCTCGGAGGAACGAAACCCTTTGAAAAGGTTCCTTTCATTTATATCGCCTCCATATGGTGCATCGTGTTTGT
TTCCTGCTGTTTCTTGTAAAACAAGTGTGGACATTCATTTAATATTTTTTCGTTATATTTTTTTGGTGACATCCTTTCTAATGCCTTATTAACCATC
GCCTGAGACCCACAGCCCTGTAGATTTCTGTGATGTTTCGGTTGCGTATTCCATAATTTTAAGCGTTTCACTTCTATTTTTTTTCATTCCTTTGAAT
TTGGATCTTAAAAAAAAAAAAAAAAAAAAAAAAAAAAAAAAAAAAAAAAAAAAAAAAAAAAAAAAAAAAAAAAAAAAAAAAA
AAAAAAAAAAAAAAAAAAAAAAAAAAAAAAAAAAAAAAAAAAAAAAAAAAA

**Note:** Remaining amplicons were not GPEET
